# Chorionic Gonadotropin Beta 7 is a marker of immune evasion in cancer

**DOI:** 10.1101/2025.05.28.656535

**Authors:** Siegen A. McKellar, Jose Mario Bello Pineda, Rohan Lattupally, Amy S. Codd, Evan W. Newell, Sydney X. Lu, Robert K. Bradley

**Affiliations:** Computational Biology Program, Public Health Sciences Division, Fred Hutchinson Cancer Center, Seattle, WA, USA; Basic Sciences Division, Fred Hutchinson Cancer Center, Seattle, WA, USA; Medical Scientist Training Program, University of Washington, Seattle, WA, USA; Molecular and Cellular Biology Graduate Program, University of Washington, Seattle, WA, USA; Department of Genome Sciences, University of Washington, Seattle, WA, USA; Division of Hematology, Stanford Medical School, Palo Alto, CA, USA; Vaccine and Infectious Disease Division, Fred Hutchinson Cancer Center, Seattle, WA, USA

**Keywords:** CGB7, chorionic gonadotropin beta, beta-hCG, immune checkpoint inhibition, immune evasion, metastatic cancer

## Abstract

Human chorionic gonadotropin beta (beta-hCG) is an oncofetal antigen expressed by trophoblast cells of the placenta, with minimal expression in adult somatic tissues. Numerous studies have demonstrated that beta-hCG-encoding genes are expressed in various cancers, but expression of these genes (*CGB3*, *CGB5*, *CGB7*, and *CGB8*) across diverse cancers has not been systematically evaluated. Here, we report that CGB genes are more widely expressed across diverse cancer types than previously appreciated and that secreted beta-hCG is readily detected. In particular, CGB genes are expressed in the majority of urothelial bladder cancers, where *CGB7* is most frequently expressed and significantly associated with an immunosuppressed tumor microenvironment, including decreased CD8^+^ T cell infiltration. Multiple CGB genes are associated with failure to respond to immune checkpoint inhibitor (ICI) therapy, and *CGB7* is particularly strongly predictive of poor prognosis. Overall, our findings indicate that beta-hCG is a clinically accessible, predictive biomarker of immunotherapeutic response.

## INTRODUCTION

Oncofetal antigens, or proteins that are canonically expressed in immune-permissive developmental contexts and are re-expressed in cancer, have garnered interest for decades as both tumor-specific biomarkers and attractive targets for anti-cancer therapeutics. Notable examples of clinically utilized oncofetal proteins include carcinoembryonic antigen (CEA) and alpha-fetoprotein (AFP). CEA is elevated in multiple malignancies and is commonly used as a biomarker of colorectal cancer, while AFP is a diagnostic marker of testicular cancer and hepatocellular carcinoma (Lazaro-Gorines et al., 2019; Terentiev et al., 2013). Due to the tumor specificity of these markers, they are both being actively investigated as a target antigen for cell therapies such as AFP-targeted CAR-T cell therapies (Sandberg et al., 2022; Munson et al., 2022).

Human Chorionic Gonadotropin Beta (beta-hCG), the beta subunit of essential heterodimeric developmental glycohormone human Chorionic Gonadotropin (hCG), is an oncofetal antigen expressed primarily in trophoblast cells during embryogenesis and reexpressed in multiple cancer types (Lapthorn et al., 1994; Douglas et al., 2014; Iles 2007). Detection of beta-hCG immunoreactivity by enzyme linked immunosorbent assay (ELISA) is the basis of modern clinical tests for pregnancy and trophoblastic disease. Elevated beta-hCG has also been observed in both trophoblastic and non-trophoblastic cancers including germ cell tumors such as choriocarcinoma, urothelial bladder cancer, lung cancer, head and neck cancer, breast cancer, cervical cancer, ovarian cancer, colorectal cancer, endometrial cancer, renal cancer, prostate cancer, and pancreatic cancer (Douglas et al., 2014; Iles 2007). Highlighting the extensive and cancer-specific expression pattern of genes encoding beta-hCG, Chew et al. developed a cancer-specificity scoring metric to screen for cancer-specific genes expressed across multiple cancer types and identified *CGB5,* one of the genes encoding beta-hCG proteins, as the gene exhibiting the strongest pan-cancer signal (Chew et al., 2019).

Beta-hCG expression and the presence of trophoblast-like cells on histology are hallmarks of aberrant trophoblastic differentiation, which can occur in non-choriocarcinomas such as urothelial bladder cancer. Trophoblastic differentiation is associated with poor prognosis and therapeutic resistance (Chang et al., 2024; Cheng et al., 2021). beta-hCG expression alone in many cancer types is associated with poor prognosis (Iles 2007).

hCG is a hormone composed of an alpha subunit encoded by *CGA* and a beta subunit encoded by one of four highly homologous genes: *CGB3* (also known as *CGB*), *CGB5*, *CGB7*, and *CGB8*.

Functioning at the maternal-fetal interface, it is essential in decidualization and the maintenance of early pregnancy via both endocrine and paracrine signaling mechanisms. hCG carries out endocrine effects as a driver of progesterone production and exerts paracrine effects that facilitate invasion (Fluhr et al., 2008; Guibourdenche et al., 2010), angiogenesis (Zygmunt et al., 2002; Berndt et al., 2006; Berndt et al., 2009), and maternal immunotolerance (Schumacher et al., 2009; Schumacher et al., 2013; Poloski et al., 2016; Dauven et al., 2016; Schumacher et al., 2017). hCG belongs to the gonadotropin family of hormones, which includes thyroid stimulating hormone (TSH), follicle stimulating hormone (FSH), and luteinizing hormone (LH). This hormone family shares a common alpha subunit, CGA. hCG, TSH, FSH, and LH have essential, but distinct, functions in pregnancy, metabolism, and in development and fertility respectively, with functional specificity conferred by unique beta subunits that possess varying degrees of homology. It is intriguing that upregulation of these specificity-conferring genes encoding beta-hCG proteins is observed in cancer.

Beta-hCG proteins are encoded by four highly homologous primate-specific coding genes arranged in tandem on chromosome 19: *CGB3, CGB5*, *CGB7*, and *CGB8*. This is in contrast with *CGB1* and *CGB2*, which have historically been classified as pseudogenes due to the presence of frameshifts that disrupt their open reading frames. *CGB3*, *CGB5*, and *CGB8* encode identical proteins classified as type 2 CGB proteins. The CGB7 protein differs by three amino acids and encodes a type 1 CGB protein. *CGB7* expression is also observed in some extra-trophoblast tissues and is uniquely regulated (Bellet et al., 1997; Sohr et al., 2011; Zimmermann et al., 2012). We refer to this collection of genes as CGB genes.

As detection and level of CGB protein is currently clinically utilized for pregnancy testing and as a clinical-grade biomarker of trophoblastic disease and germ cell tumors, we sought to explore the potential of CGB proteins as cancer biomarkers. While numerous reports suggest that expression of CGB genes may be valuable cancer biomarkers and prognostic factors, the extent to which each CGB gene is expressed in diverse cancers has not yet been systematically evaluated using the large cohorts of cancer patients available today. Furthermore, the quantitative prognostic utility of CGB expression and the specific cancer phenotypes with which CGB expression is associated remain incompletely understood, particularly across diverse cancer types.

Here, we systematically assess expression of CGB genes across diverse cancer types and reveal that all CGB genes are pervasively expressed across many cancer types, including urothelial cancer, with *CGB7* most frequently expressed. We confirm that CGB genes are transcribed and proteins are detectable as secreted proteins from cancer cells by isoform-specific qRT-PCR and bulk CGB ELISA. We determine that *CGB7* expression is strongly associated with hallmarks of an immunosuppressed tumor microenvironment by re-analyzing RNA-sequencing data, CD8+ T cell infiltration data as measured by immunohistochemistry (IHC), and tumor subtyping data from a cohort of advanced urothelial carcinoma patients. Kaplan-Meier survival analyses elucidate an association between *CGB7* expression and decreased survival probability in a cohort of advanced urothelial carcinoma patients treated with PD-L1 inhibition immunotherapy, which is supported by an association between *CGB7* expression and progressive disease in this cohort. Finally, we propose *CGB7* as a prognostic marker with strong predictive value for response to immune checkpoint inhibition via a random survival forest machine learning model of patient survival.

## RESULTS

### CGB genes are expressed in almost all cancer types

We sought to systematically evaluate the expression patterns of each functional CGB gene (*CGB3, CGB5, CGB7,* and *CGB8*) across 20 distinct cancer types utilizing gene expression data from The Cancer Genome Atlas (TCGA) **(Figure 1A)**. We confined this analysis to the 20 tumor types with at least one peritumoral control tissue sample in order to be able to accurately assess cancer specificity of CGB expression. Comparing expression of each CGB gene in tumor and matched normal peritumoral tissues across these datasets elucidated extensive cancer-specific expression of *CGB3, CGB5, CGB7,* and *CGB8* across cancer types **(Figure 1B**, **Figure 1C, Figure S1D-K)**.

**Figure 1:**
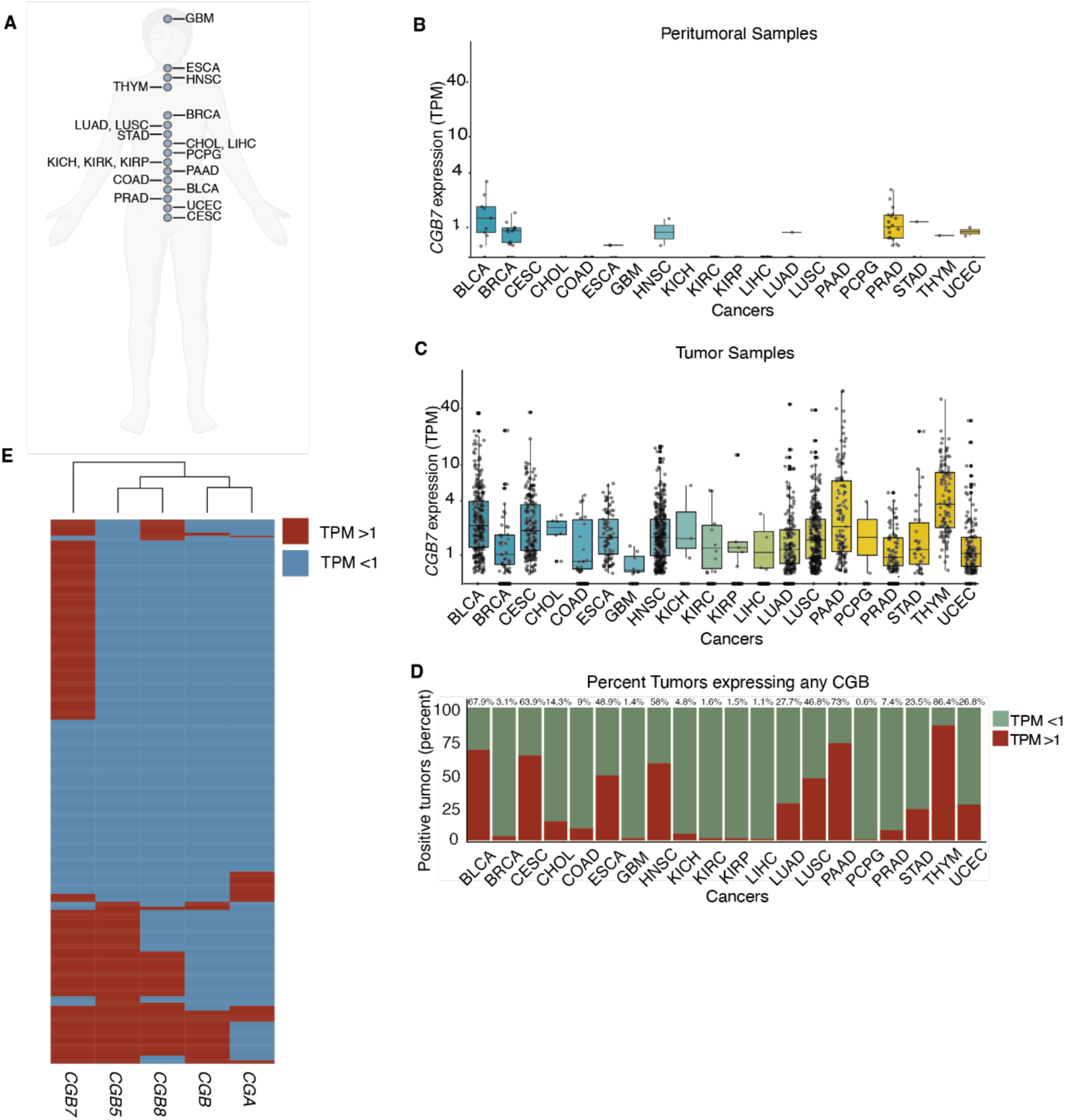
CGB is expressed in multiple cancer types. (**A**) selection of 20 unique cancer mRNA sequencing datasets from The Cancer Genome Atlas (TCGA). *CGB7* mRNA expression in healthy peritumoral tissue samples (**B**), and matched tumor tissue samples (**C**) across 20 cancer types. Cancer type reflects the site of the primary tumor. (**D**) Fraction of tumors across 20 cancer types expressing at least one CGB gene with TPM >1. Percent positive is annotated above each cancer type. (**E**) Expression status of *CGB3*, *CGB5*, *CGB7*, *CGB8*, and *CGA* in tumors from the TCGA bladder cancer (BLCA) dataset. Expressed genes are defined as TPM > 1, non-expressed genes are defined as TPM < 1. TPM = transcripts per million. Data re-analyzed from the TCGA.

Our results both confirm and notably extend previous reports of CGB expression in cancers as well as identify CGB expression in tumor types in which it has not been previously reported. For example, Cook et al. reports mRNA expression of CGB genes in 30% (n=70) of bladder cancers (Cook 2000). Hotakainen et al. reports mRNA expression of CGB genes in 50% (n=84) of the transitional cell carcinoma cases and in none of the healthy controls assayed (n=15) (Hotakainen 2002). Our new analyses reveal that nearly 68% of such tumors express one or more CGB genes at the mRNA level, while our analyses also reveal common CGB expression in tumor types such as thymoma, which has not been previously reported to our knowledge. Conversely, though *CGA* is expressed across numerous cancers, it is also expressed more frequently in numerous normal tissues, and thus expression of *CGA* does not share a cancer specific expression profile **(Figure S1A-B)**. This is consistent with the pleiotropy of *CGA*, where CGA is a subunit of all gonadotropin hormones, including thyroid stimulating hormone (TSH), follicle stimulating hormone (FSH), and luteinizing hormone (LH), which are expressed outside of development in healthy somatic tissues. Together, these data highlight the specific upregulation of CGB genes in cancer. We conclude that CGB genes are specifically and pervasively expressed in cancers and hypothesize that this specific upregulation reflects a potential novel function of CGB proteins in cancer.

We next identified cancer types that exhibited particularly high frequencies of expression of at least one CGB gene. The three cancer types with highest expression of any CGB are thymoma, pancreatic adenocarcinoma, and urothelial carcinoma **(Figure 1D)**. Trophoblastic differentiation and beta-hCG expression in urothelial cancer have been recognized for decades (Moutzouris 1993, Dirnhofer 1998, Przybycin 2020, Cheng 2021). As urothelial cancer exhibits low frequency of *CGA* expression in our analyses despite the high frequency of expression of CGB genes–highlighting the specificity of CGB gene expression in this cancer type–we selected urothelial cancer as a model to further explore the effects of CGB expression on cancer **(Figure 1D, S1A)**.

Of the four CGB genes, CGB7 is expressed most frequently across cancer types and in urothelial cancer **(Figure S2, Figure S1J, Figure S1K, Figure S1L, Figure S1M)**. We therefore sought to further explore the expression pattern of CGB7 in urothelial carcinoma. We first wondered how frequently *CGB7* is coexpressed with *CGB3*, *CGB5*, *CGB8*, and *CGA* in urothelial carcinoma.

Further evaluation of expression of these genes in the TCGA urothelial carcinoma cohort revealed that while tumors tend to co-express *CGB3*, *CGB5*, and *CGB8, CGB7*-expressing tumors cluster independently, suggesting that *CGB7* is less frequently co-expressed with *CGB3, CGB5,* and *CGB8* **(Figure 1E)**. We conclude that *CGB7* exhibits a unique expression pattern in urothelial cancer and is expressed more frequently than *CGB3, CGB5,* and *CGB8* in this dataset **(Figure 1E)**. This is consistent with a previous report of type I CGBs being the predominantly expressed type at the mRNA level in renal cell carcinoma samples (Hotakainen et al., 2006).

Furthermore, *CGB7* is frequently expressed independently of *CGA*, suggesting a potential beta-subunit specific effect independent of the hCG heterodimer **(Figure 1E)**. These findings regarding the unique expression pattern of *CGB7* are particularly notable as *CGB7* differs from *CGB3, CGB5,* and *CGB8* in sequence, tissue expression profile, and the sequences of its untranslated regulatory regions (Bellet et al., 1997; Sohr et al., 2011; Zimmermann et al., 2012).

### CGB proteins are secreted by urothelial cancer cells

The comprehensive RNA-seq data provided by the TCGA allows for systematic analysis of CGB gene expression, but these data do not directly address the question of whether expressed CGB mRNA is translated into protein. We therefore directly tested whether CGB protein is produced and secreted by urothelial cancer. We focused on urothelial cancer given the particularly high and frequent expression of CGB that we observed in this tumor type.

We first sought to validate that CGB genes are expressed in human urothelial cancer cells with an orthogonal assay and confirmed that this expression can be reversed by gene silencing. We utilized gene expression data from the Cancer Cell Line Encyclopedia (CCLE) to stratify human urothelial cancer cell lines by CGB expression **(Supplemental Table 1)**. As in human tumor samples, urothelial cancer cell lines exhibit a range of expression of CGB genes **(Table 1).**

**Table 1:** Expression of Chorionic Gonadotropin genes in cell lines from the Cancer Cell Line Encyclopedia.

| Cell Line | *CGB3* | *CGB5* | *CGB7* | *CGB8* | *CGA* | Sum of CGB genes |
| --- | --- | --- | --- | --- | --- | --- |
| KU-19-19 | 0.56 | 229.23 | 2.45 | 14.12 | 0.00 | 246.35 |
| J82 | 2.50 | 25.04 | 4.17 | 36.35 | 0.75 | 68.06 |
| UM-UC-3 | 0.36 | 0.71 | 0.41 | 28.27 | 0.00 | 29.76 |
| SCaBER | 1.42 | 2.63 | 2.20 | 16.69 | 0.45 | 22.93 |
| RT-112 | 0.28 | 3.11 | 3.79 | 10.81 | 0.33 | 18.00 |
| UM-UC-1 | 0.25 | 2.19 | 1.81 | 11.97 | 0.42 | 16.22 |
| 639-V | 0.75 | 7.79 | 1.66 | 3.78 | 0.26 | 13.98 |
| VM-CUBI | 0.50 | 3.23 | 7.22 | 1.81 | 0.00 | 12.75 |
| JMSU-1 | 0.14 | 1.36 | 1.33 | 4.37 | 0.23 | 7.19 |
| 5637 | 0.11 | 1.41 | 2.10 | 2.63 | 0.39 | 6.25 |
| HT-1197 | 0.62 | 0.45 | 2.45 | 1.98 | 1.06 | 5.50 |
| SW-1710 | 0.12 | 0.15 | 1.95 | 2.74 | 0.32 | 4.96 |
| KMBC-2 | 0.10 | 1.56 | 1.93 | 1.35 | 0.76 | 4.95 |
| 647-V | 0.11 | 1.22 | 0.37 | 1.89 | 0.00 | 3.59 |
| CAL-29 | 0.18 | 1.29 | 1.04 | 0.64 | 0.42 | 3.14 |
| HT-1376 | 0.38 | 0.74 | 1.00 | 0.29 | 9.73 | 2.41 |
| RT4 | 0.08 | 0.33 | 1.91 | 0.05 | 0.22 | 2.38 |
| BFTC-905 | 0.10 | 0.41 | 1.46 | 0.14 | 0.32 | 2.11 |
| BC-3C | 0.11 | 0.21 | 0.11 | 0.79 | 0.34 | 1.22 |
| T24 | 0.05 | 0.09 | 0.53 | 0.05 | 0.00 | 0.72 |
| TCCSUP | 0.06 | 0.08 | 0.39 | 0.08 | 0.27 | 0.62 |

Values in Transcripts Per Million (TPM).

**Table 2:** Cox Proportional Hazards Regression for Overall Survival.

| Characteristic | Univariate |  |  |  | Multivariate<sup>a,b</sup> |  |  |  |
| --- | --- | --- | --- | --- | --- | --- | --- | --- |
|  | HR | 95% CI | p-value | q-value<sup>c</sup> | HR | 95%CI | p-value | q-value<sup>c</sup> |
| **Tumor mutational burden<sup>d</sup>** | 0.43 | 0.28 - 0.66 | 1.10 x 10<sup>-4</sup> | 7.72 x 10<sup>-4</sup> | 0.32 | 0.19 - 0.50 | 1.10 x 10<sup>-6</sup> | 7.73 x 10<sup>-6</sup> |
| ***CGB7* expression<sup>e</sup>** | 1.44 | 1.09 - 1.89 | 9.05 x 10<sup>-3</sup> | 0.04 | 1.48 | 1.05 - 2.11 | 0.026 | 0.093 |
| **ECOG Performance Status<sup>f</sup>:** |  |  |  |  |  |  |  |  |
| 0 (reference) | - | - | - | - | - | - | - | - |
| 1 | 2.1 | 1.58 - 2.79 | 2.95 x 10<sup>-7</sup> | 4.14 x 10<sup>-6</sup> | 2.62 | 1.85 - 3.71 | 6.07 x 10<sup>-8</sup> | 8.50 x 10<sup>-7</sup> |
| 2 | 1.97 | 1.04 - 3.73 | 0.036 | 0.13 | 3.32 | 1.53 - 7.17 | 2.30 x 10<sup>-3</sup> | 0.011 |
| **Received platinum:** |  |  |  |  |  |  |  |  |
| No (reference) | - | - | - | - | - | - | - | - |
| Yes | 1.41 | 1.01 - 1.98 | 0.047 | 0.13 | 1.54 | 1.03 - 2.30 | 0.037 | 0.1 |
| **Tumor cell PD-L1<sup>g</sup>:** |  |  |  |  |  |  |  |  |
| < 1% (reference) | - | - | - | - | - | - | - | - |
| ≥ 1% but < 50% | 1.12 | 0.66 - 1.90 | 0.66 | 0.99 | 1.14 | 0.64 - 2.02 | 0.65 | 0.83 |
| ≥ 50% | 1 | 0.70 - 1.44 | 0.99 | 0.99 | 0.77 | 0.49 - 1.23 | 0.27 | 0.54 |
| **Intravesical BCG administered:** |  |  |  |  |  |  |  |  |
| No (reference) | - | - | - | - | - | - | - | - |
| Yes | 0.98 | 0.73 - 1.32 | 0.89 | 0.99 | 0.92 | 0.63 - 1.33 | 0.65 | 0.83 |
| **Tobacco use history:** |  |  |  |  |  |  |  |  |
| Never (reference) | - | - | - | - | - | - | - | - |
| Previous | 0.9 | 0.68 - 1.19 | 0.45 | 0.78 | 1.01 | 0.72 - 1.42 | 0.95 | 0.95 |
| Current | 1.02 | 0.65 - 1.61 | 0.92 | 0.99 | 1.09 | 0.63 - 1.90 | 0.75 | 0.87 |
| **Stage:** |  |  |  |  |  |  |  |  |
| I (reference) | - | - | - | - | - | - | - | - |
| II | 0.96 | 0.69 - 1.35 | 0.83 | 0.99 | 1.25 | 0.80 - 1.95 | 0.33 | 0.57 |
| III | 1.23 | 0.86 - 1.75 | 0.25 | 0.51 | 1.4 | 0.90 - 2.19 | 0.14 | 0.32 |

| IV | 0.98 | 0.68 - 1.42 | 0.92 | 0.99 | 0.89 | 0.54 - 1.46 | 0.65 | 0.83 |
| --- | --- | --- | --- | --- | --- | --- | --- | --- |
| **Sex:** |  |  |  |  |  |  |  |  |
| **Male**<br>**(reference)** | - | - | - | - | - | - | - | - |
| **Female** | **1.23** | **0.91 - 1.66** | **0.18** | **0.42** | **1.05** | **0.72 - 1.53** | **0.81** | **0.87** |

<sup>a</sup>Log-rank test: p-value = 1.05 x 10<sup>-7</sup>
<sup>b</sup>Akaike Information Criterion =1736.51; Bayesian Information Criterion = 1780.66; Harrell's Concordance Index = 0.67
<sup>c</sup>Benjamini-Hochberg FDR correction
<sup>d</sup>log<sub>10</sub>(number of missense mutations)
<sup>e</sup>log<sub>10</sub>[CGB7 expression (TPM)]
<sup>f</sup>Eastern Oncology Cooperative Group Performance Status
<sup>g</sup>Fraction of tumor cells with positive PD-L1 staining

Interestingly, *CGB3*, *CGB5,* and *CGB8* frequently exhibited markedly higher expression than did *CGB7* in these cell lines – in contrast to the primary tumor mRNA expression data from the TCGA, where all CGB genes were expressed at a similar level (**Table 1**). However, this observation is consistent with prior findings that type II CGBs (CGB3, CGB5, and CGB8) exhibited higher expression than type I CGBs (CGB7) in urothelial cancer (Hotakainen et al., 2007). *CGA* expression was low or absent from the majority of these cell lines (**Table 1**).

We selected seven urothelial cancer cell lines with high summed CGB expression and one cell line with undetectable CGB expression as examples of CGB-positive and CGB-negative tumors to validate CGB gene expression and protein levels *in vitro*. All cell lines selected are negative for *CGA* expression at the mRNA level. We observed expression of *CGB3, CGB5,* and *CGB8* across the seven positive cell lines surveyed, but not in the negative cell line **(Figure 2A)**. Treatment of J82 cells with a pan-CGB targeting siRNA pool depleted signal from the CGB3/5/8-specific probe in J82 cells **(Figure 2B)**, confirming that the detected mRNA signal arises from expression of CGB genes. We observed clear but low *CGB7* expression in one of the seven assayed cell lines (SCaBER cells; **Figure 2C)**. *CGB7* expression is too low to be reliably detected in the remaining cell lines not shown. This is supported by the lower expression of *CGB7* relative to other CGB genes that we observed in gene expression data from CCLE data **(Table 1)**.

**Figure 2:**
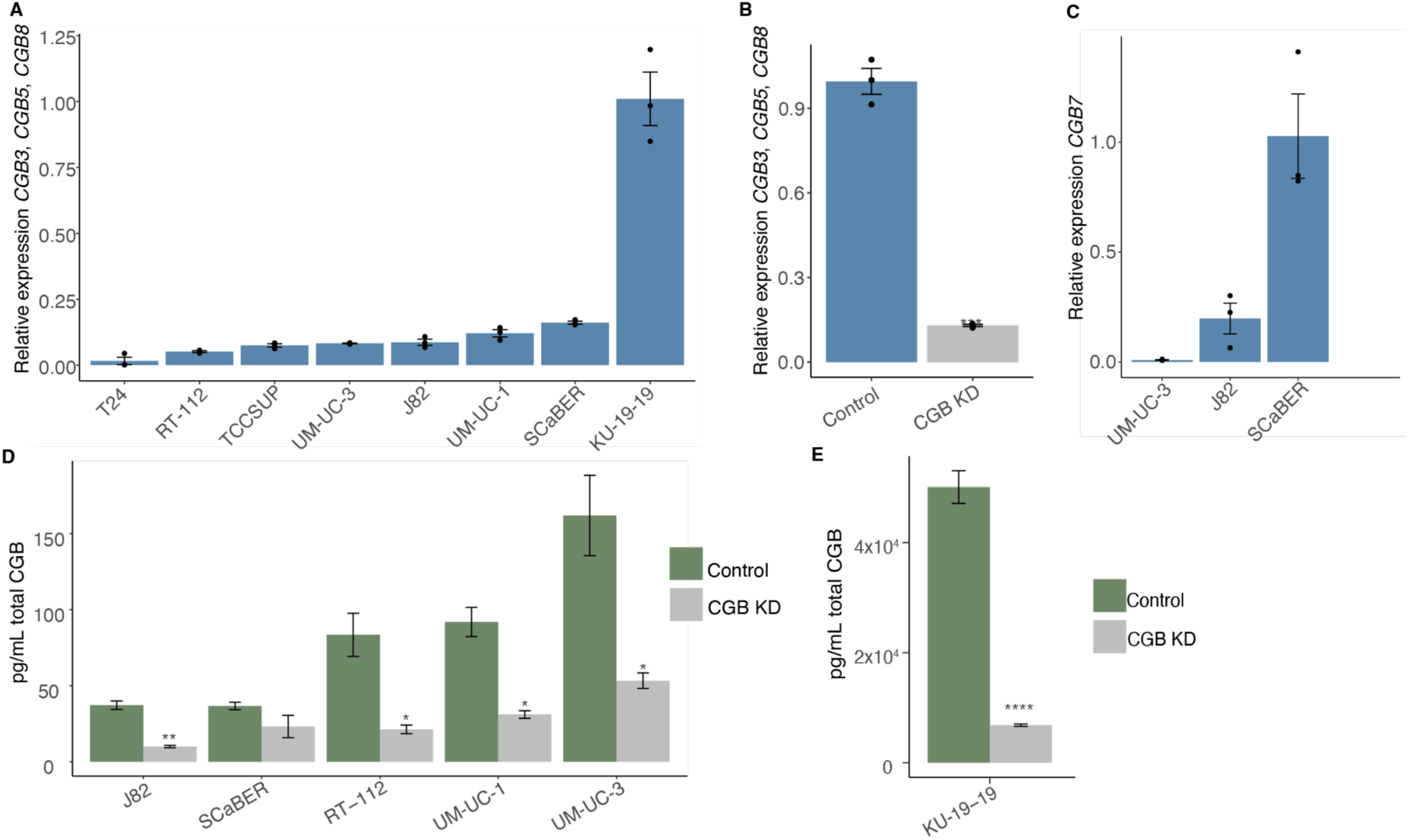
**CGB is expressed in multiple human urothelial carcinoma cell lines.** (**A**) Expression of bulk *CGB3*, *CGB5*, and *CGB8* mRNA normalized to *GAPDH* quantified by RT-qPCR in human urothelial carcinoma cell lines. Expression is relative to the cell line with the highest expression. Data in Figure 2A–Source Data 2A file. (**B**) Expression of *CGB3*, *CGB5*, and *CGB8* mRNA normalized to *GAPDH* quantified by RT-qPCR in J82 cells treated with a non-targeting siRNA pool or an siRNA pool targeting all CGB genes. Expression is relative to the cell line with the highest expression. P-value calculated via welch two sample t-test. Data in Figure 2B–Source Data 2B file. (**C**) Expression of *CGB7* mRNA normalized to *GAPDH* quantified by RT-qPCR in human urothelial carcinoma cell lines. Expression is relative to the cell line with the highest expression. (**D**) Secretion of total CGB protein quantified by ELISA in human urothelial carcinoma cell lines treated with a non-targeting siRNA pool or an siRNA pool targeting all CGB genes. pg/mL CGB protein shown per 10,000 cells. P-value calculated via welch two sample t-test. Data in Figure 2D–Source Data 2D file. (**E**) Secretion of total CGB protein quantified by ELISA in KU-19-19 cells treated with a non-targeting siRNA pool or an siRNA pool targeting all CGB genes. pg/mL CGB protein shown per 10,000 cells. P-value calculated via welch two sample t-test. All experiments run in biological and technical triplicate; all error bars represent SE; *p < 0.5; **p < 0.01; ****p < 0.0001.

We next validated that CGB proteins are translated and secreted as free beta subunits and that the signal arises from CGB mRNA. Previously, secretion of CGB proteins from several urothelial cancer cell lines was shown (Iles et al., 1987). Iles et al. did not confirm reversibility of signal with knockdown of CGB genes, so we sought to confirm specificity of detection with pan-CGB gene knockdown. We selected J82, SCaBER, RT-112, UM-UC-1, UM-UC-3, and KU-19-19 cells given both their high expression of CGB genes and absent *CGA* expression by mRNA analysis and performed ELISA to detect total CGB protein from supernatant **(Figure 2D, 2E)**.

These data confirmed that CGB protein is readily produced and detectable as a secreted protein from all cell lines, with KU-19-19 cells producing particularly notable amounts. As *CGA* is not appreciably expressed in these cell lines **(Table 1)**, we infer that CGB proteins are stable and secreted unbound to CGA. Specificity of the assay was further supported by targeted knock down of bulk CGB genes, resulting in a significant decrease in detected CGB protein **(Figure 2D, 2E)**.

We conclude that both type-I-encoding (*CGB7*) and type-II-encoding (*CGB3, CGB5,* and *CGB8*) CGB genes are expressed in human urothelial cancer cells and that CGB proteins are stable as free beta subunits and are secreted from cancer cells.

### *CGB7* expression is associated with reduced anti-tumor immune activity

As *CGB7* is the most frequently expressed of the CGB genes in our analyses of the TCGA urothelial carcinoma cohort, we elected to focus on *CGB7*. To interrogate the possible functional effects of *CGB7* expression in cancer, we re-analyzed RNA-sequencing data from the TCGA primary urothelial carcinoma cohort by comparing the gene expression profiles of *CGB7*-positive and *CGB7*-negative tumors. Gene Ontology (GO) enrichment analysis revealed that downregulated genes in *CGB7*-positive tumors are enriched for diverse biological processes **(Figure S3A)**. Interestingly, one of the highest-ranked enriched GO terms was related to immune response, which we found to be particularly intriguing given previous literature demonstrating a functional role for hCG in immune tolerance in pregnancy (Schumacher et al., 2009; Schumacher et al., 2013; Poloskiet al., 2016; Dauven et al., 2016; Schumacher et al., 2017).

Further probing of associated GO terms revealed that *CGB7* expression was specifically associated with downregulation of genes involved in the adaptive immune response, lymphocyte activation, and positive immune system regulation, indicating that *CGB7* expression is inversely associated with inflammation **(Figure 3A)** in a manner reminiscent of hCG’s documented role in maternal immunotolerance.

**Figure 3:**
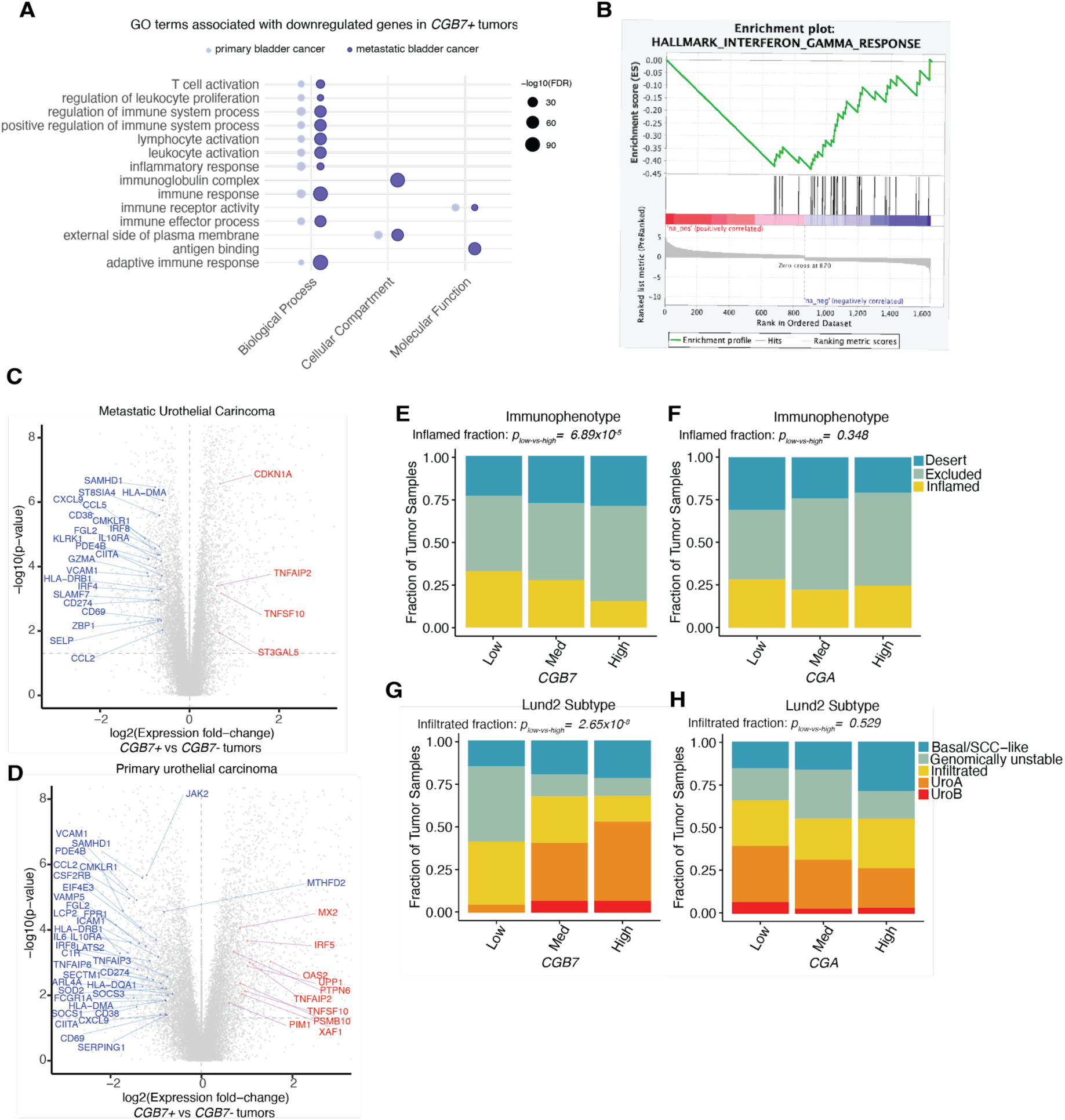
**CGB expression correlates with altered immune infiltrate.** (**A**) Immune-associated Gene Ontology terms associated with genes downregulated in *CGB7+* tumors in metastatic urothelial cancer (data re-analyzed from Mariathasan et al., 2018) and in primary urothelial cancer (data re-analyzed from TCGA BLCA cohort). *CGB7*+: >1 TPM. *CGB7*-: <0.25 TPM. (**B**) Interferon Gamma response genes are negatively correlated with *CGB7* expression in metastatic urothelial tumors (data re-analyzed from Mariathasan et al., 2018). GSEA normalized Enrichment Score (NES): −2.20. FWER p-Value: 0.004. (**C**) Differentially expressed genes in *CGB7*+ vs *CGB7*-metastatic urothelial tumors (data re-analyzed from Mariathasan et al., 2018) as determined by Mann-Whitney U Test. Interferon gamma response genes that are significantly differentially expressed are in red (upregulated) or blue (downregulated) and reach p-value < 0.05 and log fold change > 1.5. CGB7+: >1 TPM. CGB7-: <0.25 TPM. (**D**) Differentially expressed genes in *CGB7*+ vs *CGB7*-primary urothelial tumors (data re-analyzed from TCGA) as determined by Mann-Whitney U Test. Interferon gamma response genes that are significantly differentially expressed are in red (upregulated) or blue (downregulated) and reach p-value < 0.05 and log fold change > 1.5. CGB7+: >1 TPM. CGB7-: <0.25 TPM. (**E**) Immunophenotype data in tumors stratified by *CGB7* expression: immune desert, immune excluded, or inflamed. P-values determined by proportions test with continuity correction. Chi-squared = 15.84, df = 1, p-value = 6.89×10^-5^. (**F**) As in (**E**), but stratified by *CGA* expression. Chi-squared = 0.88, df = 1, p-value = 0.35. (**G**) Tumor subtype data in tumors stratified by *CGB7* expression: basal/SCC-like, genomically unstable, immune infiltrated, urothelial type A, or urothelial type B. P-values determined by proportions test with continuity correction. Chi-squared = 30.95, df = 1, p-value = 2.65×10^-8^. (**H**) As in (**G**), but stratified by *CGA* expression. Chi-squared = 0.40, df = 1, p-value = 0.53. Med = medium.

To confirm this finding in an orthogonal dataset, we analyzed mRNA-sequencing data from the IMVigor 210 multicentre, single-arm, phase II clinical trial (Mariathasan et al., 2018). This cohort is composed of patients with advanced urothelial carcinoma that received Atezolizumab, an anti-PD-L1 checkpoint inhibitor, and is ideally suited for our investigation given that pre-treatment samples were characterized by both RNA-seq and immunohistochemistry (Mariathasan et al., 2018). We again found that genes downregulated with *CGB7* expression were strongly associated with inflammatory processes **(Figure 3A)**. Notably, in genes that are downregulated in *CGB7*-positive tumors, at least 17 of the 20 highest ranked enriched GO terms are robustly associated with inflammation and immunity, reinforcing a negative association between *CGB7* and inflammation **(Figure S3B)**. The association between *CGB7* expression and regulation of immune response in this dataset is further supported by pronounced downregulation of genes involved in interferon gamma response in *CGB7*+ tumors identified by gene set enrichment analysis via GSEA software (Subramanian & Tamayo et al., 2005; Mootha & Lindgren et al., 2003). **(Figure 3B)**.

Interferon gamma signaling is strongly associated with response to immune checkpoint inhibition, and indeed, disruption of interferon gamma signaling remains a mechanism of resistance to ICI (Zaretsky et al., 2016; Grasso et al., 2020). Exploring this *CGB7*-associated downregulation of interferon gamma response genes in both the IMVigor210 cohort dataset and in the TCGA urothelial carcinoma cohort dataset reveals downregulation of numerous genes encoding inflammatory cytokines and chemokines, immune checkpoints such as PD-L1, and HLA proteins involved in antigen presentation; all features of immune evasion in cancer **(Figure 3C, 3D)**. As hCG is well-established as a facilitator of maternal immunosuppression (Schumacher et al., 2009; Schumacher et al., 2013; Poloski et al., 2016; Dauven et al., 2016; Schumacher et al., 2017), we hypothesize that *CGB7* may similarly exhibit immunosuppressive functions that ultimately promote immune escape of tumors.

As interferon gamma facilitates tumor infiltration of CD8+ T cells and is ultimately a marker of CD8+ T cell activity, we next investigated whether *CGB7* expression affects CD8+ T cell infiltration using immune phenotype classification data for tumor samples from the IMVigor 210 cohort. Here, immunophenotype is determined by CD8a IHC staining of FFPE tumor sections (Mariathasan et al., 2018). Tumors were scored on the basis of CD8a staining frequency and pattern as exhibiting an immune infiltrated, immune excluded, or immune desert phenotype (Mariathasan et al., 2018). We stratified tumors by *CGB7* expression and binned samples according to negative-low (<25%), moderate (25%-75%), or high (>75%) *CGB7* expression.

Comparing the distribution of immunophenotypes between *CGB7*-low and *CGB7*-high tumors elucidates a significantly distinct distribution of inflamed tumors (p=6.89×10^-5^) **(Figure 3E)**. We observed no statistically significant differences in the distributions of immune excluded or immune desert phenotypes between *CGB7*-low and *CGB7*-high tumors, though there is a trend toward higher incidence of immune exclusion in *CGB7*-high tumors (p= 0.062, p= 0.32). These findings that *CGB7*-high tumors exhibit decreased immune infiltration is consistent with our findings that *CGB7* expression is associated with downregulated IFNg signaling, as IFNg is predominantly secreted by activated lymphocytes, promotes tumor infiltration of immune cells such as CD8+ T cells, and is a marker of ICI response (Grasso et al., 2020).

Furthermore, when we compare *CGA*-low and *CGA*-high tumors, there is no significant difference in the distribution of inflamed tumors (p=0.35), supporting a possible *CGB7*-specific effect on CD8+ T cell exclusion **(Figure 3F)**.

Additionally, when we explored associations between *CGB7* expression and subtype as scored by Mariathasan et al. according to the Lund classification system, an established system for urothelial cancer classification (Sjödahl et al., 2012; Sjödahl et al., 2013), we observe a significantly decreased fraction of immune-infiltrated tumors in *CGB7*-high tumors compared with *CGB7*-low tumors (p=2.65×10^-8^) **(Figure 3G)**. This infiltrated class of tumors is described as enriched for immune system processes and lymphocyte activation, including an enrichment of activated T cell markers and cytotoxic T cell effector genes (Sjödahl et al., 2012). There was no association between the immune infiltrated subtype and *CGA* expression, further supporting an association between *CGB7* expression and reduced immune infiltration (p=0.53) **(Figure 3H)**.

We focused these and prior analyses on *CGB7* because *CGB7* is more commonly expressed than other CGB genes in this cancer type **(Figure 1E)**. Because some CGB-positive samples express only genes encoding type I CGBs (*CGB7*), others express only genes encoding type II CGBs *(CGB3, CGB5,* and *CGB8*), and others both type I and type II, we sought to test whether the functional association between CGB gene expression and immunophenotypes was likely driven by genes encoding type I or type II CGBs (or both). We used a patient stratification approach to address this question.

We first tested whether the associations between *CGB7* and CD8+ T cell infiltration and tumor subtype are upheld independently of *CGB3, CGB5,* and *CGB8* expression. We selected samples negative for *CGB3, CGB5,* and *CGB8* expression, and to ensure sufficient samples for analysis, we lowered the threshold for negative expression from 0.25 to 1 TPM to increase sample size. We then stratified samples by negative-low, moderate, or high *CGB7* expression as described. Consistent with our prior findings, *CGB7*-high tumors exhibit significantly decreased CD8+ T cell infiltration (incidence of inflamed tumors) compared with CGB7-low tumors (p=1.60×10-6) **(Figure S3C)**. Similarly, *CGB7*-high tumors exhibit a significantly decreased fraction of immune infiltrated tumors compared with *CGB7*-low tumors (p=0.00450) **(Figure S3D)**. Both of these findings are independent of *CGB3, CGB5,* and *CGB*8 expression. Conversely, selecting *CGB7*-negative samples and stratifying by summed *CGB3, CGB5*, and *CGB8* expression, we find no association between *CGB3*, *CGB5*, and *CGB8* expression and CD8+ T cell infiltration (p=0.730) or incidence of immune-infiltrated tumors (p=1) independent of *CGB7* expression **(Figure S3E, S3F)**. It is important to note, however, that our statistical power for these analyses was not equal for type I and type II CGB analyses due to *CGB7* being the most commonly expressed CGB gene **(Figure 1E)**.

Overall, these findings reinforce our hypothesis that *CGB7* is consistently associated with an immunosuppressive tumor microenvironment and hallmarks of immune escape. We conclude that *CGB7* is significantly associated with blunted interferon gamma signaling and is specifically and significantly associated with decreased CD8+T cell infiltration in the urothelial carcinoma cohorts investigated.

### CGB expression is associated with decreased response to ICI therapy

As interferon gamma release and CD8+ T cell infiltration are associated with response to ICI therapy, we next wondered whether *CGB7* expression had implications for overall survival and response to anti-PD-L1 checkpoint inhibition. To investigate this the effect of *CGB7* on survival, we selected *CGB7*-positive (TPM >1) and *CGB7*-negative (TPM < 0.25) tumors and performed Kaplan Meier survival analysis. We found that *CGB7* expression is indeed associated with significantly decreased overall survival probability (p= 6.87×10^-3^) **(Figure 4A)**. Though this decrease in survival probability associated with *CGB7* expression is strongest, *CGB3* and *CGB5,* but not *CGB8*, are also associated with significantly decreased survival probability **(Figure S4A-C)**. In contrast, *CGA* expression had no effect on overall survival **(Figure 4B)**.

**Figure 4:**
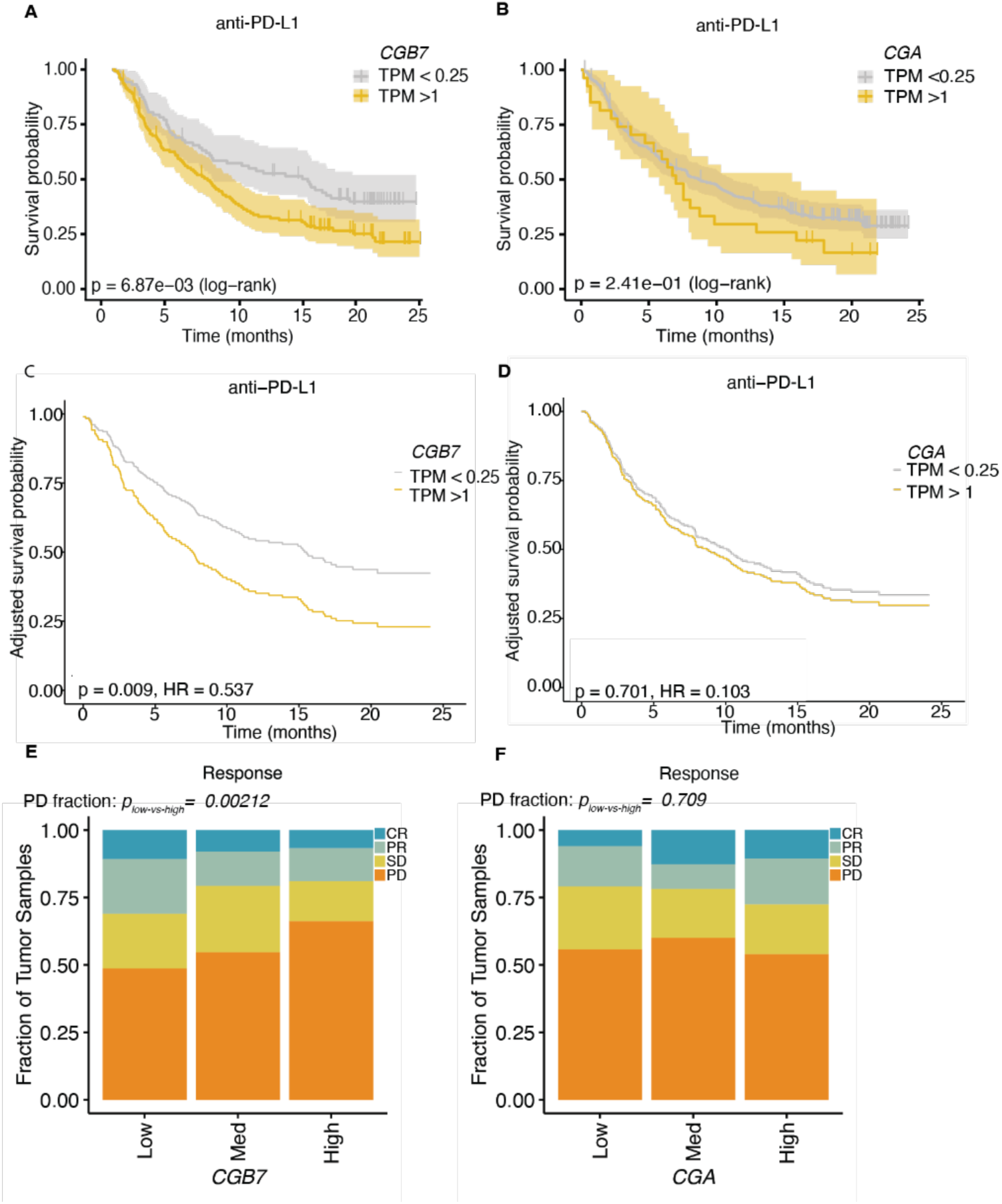
**CGB expression is associated with decreased response to ICI therapy** (**A**) Kaplan Meier overall survival curves comparing patients with *CGB7*+ (gold) and *CGB7*-(gray) advanced urothelial tumors undergoing treatment with Atezolizumab (anti-PD-L1). Data re-analyzed from Mariathasan et al., 2018. P-value determined by log rank test. The estimated survival functions (solid), censored events (crosses), and 95% confidence intervals (shaded regions) are shown. (**B**) As in (**A**), but patients stratified by *CGA* expression. (**C**) Kaplan Meier overall survival curves as shown in (**A**) adjusted for confounding effects of sex and tumor mutational burden covariates by Cox Proportional Hazards modeling. Hazard ratio (HR) and p-value obtained from fitting a Cox Proportional Hazards model. (**D**) Kaplan Meier overall survival curves as shown in (**B**) adjusted as described in (**C**). (**E**) Response determined by RECIST criteria (data re-analyzed from Balar et al., 2017). P-values determined by proportions test with continuity correction. Chi-squared = 9.44, df = 1, p-value = 0.0021. (**F**) As in (**A**), but stratified by *CGA* expression. Chi-squared = 0.14, df = 1, p-value = 0.71. Response Evaluation Criteria in Solid Tumors (RECIST) scoring: CR = complete response, PR = partial response, SD = stable disease, PD = progressive disease. Med = medium.

We wondered whether these associations between CGB gene expression and survival persist when controlling for common potential covariates tumor mutational burden (TMB) and sex. TMB is a known predictor of response to immune checkpoint inhibition (Samstein et al., 2019; Ricciuti et al., 2022; Hellmann et al., 2019; Hellmann et al., 2018). Sex is associated with differential survival outcomes in some checkpoint immunotherapy cohorts (Jang et al., 2021; Ye et al., 2020) and in this specific dataset (**Figure S4I**). Even when survival probabilities are adjusted for the contributions of TMB and sex using Cox proportional hazards modeling, the adjusted survival probability of patients with *CGB7*-positive cancers remains significantly decreased while *CGA* has no effect on adjusted overall survival probability (p= 0.009, p= 0.70) **(Figure 4C, 4D)**. Similarly, the adjusted survival probabilities of patients with *CGB3*-positive and *CGB5*-positive tumors remains significantly decreased **(Figure S4D-F)**. We conclude that *CGB3, CGB5,* and *CGB7* expression are associated with a significant reduction in survival probability in metastatic urothelial carcinoma patients receiving immune checkpoint blockade, with *CGB7* expression exhibiting the most robust association.

We next wondered whether *CGB7* expression impacts response to checkpoint immunotherapy. Each patient in the IMvigor 210 clinical trial was evaluated for response to anti-PD-L1 therapy by the Response Evaluation Criteria in Solid Tumors (RECIST) and scored as exhibiting progressive disease (PD), stable disease (SD), partial response (PR), or complete response (CR) (Balar et al., 2017). Again binning patients by *CGB7* expression corresponding to negative-low, moderate, or high expression levels, we compared the proportion of each clinical response classification in *CGB7*-high vs *CGB7*-low tumors to determine patterns of response associated with *CGB7* expression. *CGB7*-high tumors exhibited a significantly higher incidence of PD (p=0.0021), but no significant difference in incidence of CR (p = 0.25), PR (p = 0.051), or SD (p = 0.25) **(Figure 4E)**. Conversely, comparing incidence of PD between *CGA*-high vs *CGA*-low tumors demonstrated no significant differences (p=0.71). Similarly, we observed no significant differences in incidence of CR (p = 0.074), PR (p =0.53), or SD (p = 0.12) between *CGA*-high vs *CGA*-low tumors **(Figure 4F)**, highlighting a potential beta-subunit-specific effect on overall survival.

Repeating these analyses to parse *CGB7*-specific effects independent of *CGB3*, *CGB5*, and *CGB8* expression as described above, we determined that high *CGB7* expression is still associated with significantly increased incidence of progressive disease compared with low *CGB7* expression independently of *CGB3, CGB5,* and *CGB8* expression (p=0.0016) **(Figure S4G)**. Interestingly, when *CGB7*-high samples are filtered out to facilitate bulk *CGB3, CGB5,* and *CGB8* analyses independent of *CGB7* expression, we observe that *CGB3, CGB5,* and *CGB8* are also significantly associated with increased incidence of progressive disease (p=0.0027) independently of *CGB7*, indicating that general CGB expression is associated with progressive disease and thus resistance to ICIs **(Figure S4H).**

Taken together, we conclude that *CGB7* expression is strongly associated with an immunosuppressive tumor microenvironment and decreased overall survival probability, suggesting a role of *CGB7* in immune escape of tumors. We additionally conclude that *CGB3* and *CGB5* are associated with decreased survival probability. Ultimately, expression of genes encoding both type I and type II CGB genes are associated with resistance to ICIs in advanced urothelial carcinoma

### CGB expression predicts response to ICI

We next sought to analyze the effect of *CGB7* expression on survival probability by further evaluating our Cox Proportional Hazards model. We first considered the effects of clinical variables collected in the IMVigor 210 trial, including *CGB7* expression, on overall survival using univariate Cox Proportional Hazards modeling, where variables are assessed separately, and in the multivariate setting, where confounding effects of all variables are controlled for **(Figure S5A)**. In both approaches, tumor mutational burden (TMB), Eastern Cooperative Oncology Group (ECOG) status, and *CGB7* expression, which we treat as a continuous variable, were statistically significant prognostic variables associated with overall survival. Consistent with this notion, our analyses show that TMB is associated with a decreased risk of death (HR = 0.32, p = 1.10 x 10^-6^) after controlling for all other covariates. Similarly, ECOG status negatively impacts patient survival during ICI therapy as demonstrated in the original Phase II trial of the IMvigor210 study (Balar et al., 2017), which is reflected in our own analyses (ECOG 1: HR 2.62, p = 6.07 x 10^-8^; ECOG 2: HR 3.32, p = 2.30 x 10^-3^). Accordingly, increased *CGB7* expression is correlated with increased risk of death (HR = 1.48, p = 0.026) after controlling for possible confounding (**Figure S5B**).

We next aimed to quantify the predictive value of *CGB7* as a prognostic marker over time using inferences from a Random Survival Forest (RSF) model, a machine learning ensemble consisting of multiple survival trees (Ishwaran 2008). RSF modeling enables us to predict the time-to-death in our patient cohort, and has been used widely in the literature to predict the prognosis of patients in various disease and interventional contexts (Ishwaran & Malley, 2014; Pineda et al., 2024; Picket et al., 2021; Zhao et al., 2024; Sarica et al., 2023; Bohannan et al., 2022; Jung et al. 2019). To grow the RSF, we randomly selected 70% of the patients in the cohort, utilized 1000 base learners, and included the following covariates as input: ECOG status, TMB, *CGB7* expression, platinum chemotherapy history, stage, tobacco use, sex, tumor cell PD-L1 level, and intravesical BCG administration. The resultant model had an Out-of-Bag (OOB) error of 38.9%. The OOB error is calculated from predictions for unsampled patients from each bootstrapping iteration, and represents an unbiased estimate of the test error. We also directly measured model performance using the remaining 30% of patients excluded from model training, this test error measuring 35.6% (**Figure S5B**). The RSF survival curve estimates closely mirror those generated by Kaplan Meier survival analysis, lending confidence to our model (**Figure 5A**).

**Figure 5:**
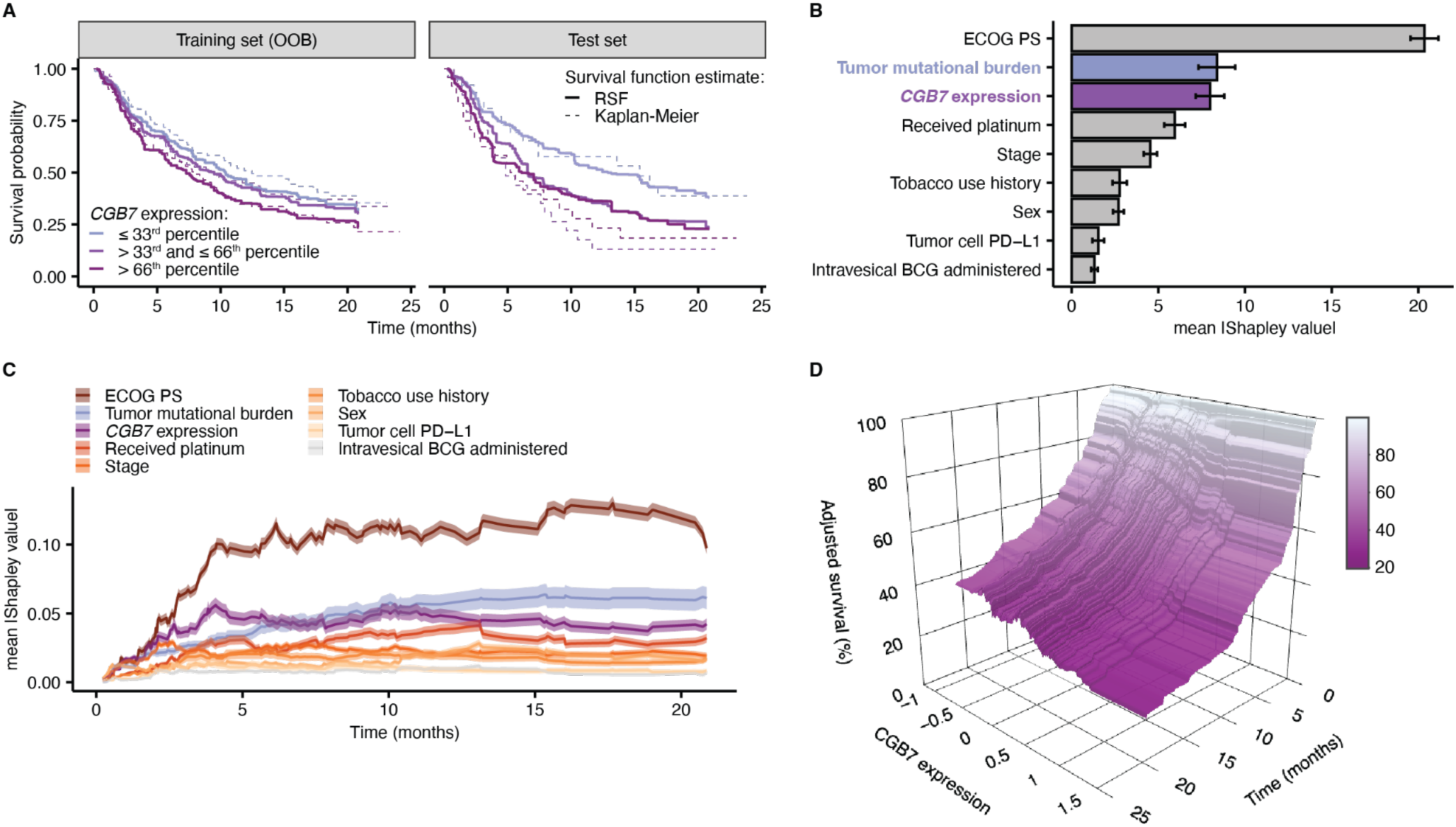
***CGB7* expression is a prognostic variable associated with decreased survival** (**A**) Survival estimates from the Random Survival Forest (RSF) model (median prediction, solid lines) and the Kaplan–Meier estimate (dashed lines) from the training (OOB) and test sets. Patients are stratified into terciles based on *CGB7* expression. (**B**) RSF feature importance quantified via the mean absolute Shapley values. Error bars denote the 95% confidence intervals. (**C**) Time-dependent Shapley values, corresponding to survival estimates at specific timepoints. The 95% confidence interval of the mean (transparent ribbon) is shown. (**D**) Adjusted survival probability measured via partial dependence, as a function of time and *CGB7* expression.

The survival probability predictions produced by the RSF model suggest that increasing *CGB7* expression is correlated with a decrease in median survival time, and conform with the aforementioned findings from the Cox Proportional Hazards regression (**Fig 5A, Fig S5C**). To quantify the contributions of *CGB7* expression to the predictions of time-to-death generated by the RSF model, we calculated the Shapley values corresponding to each variable utilized in our model (Lundberg & Lee, 2017; Maksymiuk et al., 2020; Shapley, 1953; Štrumbelj & Kononenko, 2014). Shapley values associated with the RSF prediction for mortality, the number of expected deaths over the observation window, were calculated which showed the expected protective effect of high TMB and the increased risk of death conferred by increased *CGB7* expression **(Figure S5D-E)**. Ranking Shapley values by the mean magnitude calculated over the entire cohort allows quantification of a feature’s importance to the model prediction relative to the other model covariates. In these analyses, *CGB7* expression scored highly behind ECOG performance status and equivalently to TMB (**Figure 5B**). We extended these analyses to a time-dependent implementation. First, we verified the time frame upon which inferences from the RSF model are valid, through time-dependent Receiver Operating Characteristic (ROC) analyses. We calculated the cumulative/dynamic area under the ROC curve (AUC^C/D^) which quantifies the model’s accuracy at differentiating patient deaths occurring before a particular time point, versus those who survive beyond this time. Our analyses suggest a time horizon up to ∼ 15 months (**Figure S5F**). Second, we calculated Shapley values associated with time point-specific survival probability, which suggest that *CGB7* expression has a dynamic relative contribution over time but preserved high importance throughout the observation window.

Interestingly, *CGB7* expression is linked to greater prediction contributions compared to TMB at earlier times, up to ∼ 6 months (**Figure 5C**). To obtain the adjusted or marginal effects of *CGB7* expression on overall survival, we implemented time-dependent partial dependence analyses which show a clear negative correlation between *CGB7* expression and survival probability **(Figure 5D)**. Ultimately, we observe a robust association between *CGB7* expression and predicted survival probability, in which *CGB7* expression is associated with reduced time to death in advanced urothelial cancer in the context of immune checkpoint inhibition.

## DISCUSSION

We identify CGB genes as biomarkers of immune evasion in cancer and as prognostic factors for response to immune checkpoint inhibition. We find that all CGB genes are extensively expressed across cancer types, with *CGB7* being most frequently expressed. We confirm that CGB genes are expressed and the resultant proteins secreted from urothelial carcinoma cells *in vitro*. In a cohort of advanced urothelial carcinoma patients, *CGB7* expression is associated with reduced CD8+ T cell infiltration. *CGB7* and bulk expression of *CGB3*, *CGB5,* and *CGB8* are associated with reduced response to anti-PD-L1 checkpoint immunotherapy by RECIST criteria. We subsequently identify that tumors expressing *CGB7, CGB3,* or *CGB5* are significantly associated with decreased survival probability via Kaplan-Meier survival analyses, and this association is upheld with correction for TMB and sex via Cox Proportional Hazards modeling. As *CGB7* expression is most frequently observed in urothelial carcinoma and is associated with markers of immune evasion, we focused on investigating the predictive value of *CGB7* and ultimately demonstrate the prognostic value of *CGB7* as a marker of poor prognosis in urothelial carcinoma in the context of immune checkpoint inhibition using Random Survival Forest modeling. In fact, *CGB7* is the second most important variable in survival predictions, behind TMB, early in cancer progression. Taken together, our data suggest that *CGB7* may facilitate the conversion of the tumor microenvironment to an immunosuppressive state.

One limitation of this study is the lack of functional characterization of CGB proteins in cancer, as functional immunology efforts are made challenging by the primate-specific nature of CGB genes and lack of murine orthologs. These functional studies, however, will be necessary to establish the immunosuppressive mechanism of action of CGB7. Our findings here demonstrate that, with careful experimental design and immune competent models, the study of CGB7 as an immunomodulator is ripe for further functional exploration. The primate-specificity of CGB gene expression further reinforces the value of analyzing high quality patient data, and the value of extending our analyses of *CGB7* to additional patient datasets.

A second limitation is the limited power of our current analyses dissecting isolated phenotypic associations with *CGB7* expression and with bulk summed *CGB3, CGB5,* and *CGB8* expression due to study size. In the analyses that we present here, it is interesting that, while *CGB7* is the only CGB gene associated with markers of immune evasion, multiple CGB genes are associated with ICI resistance and decreased survival. This might reflect a unique mechanism of action of *CGB7*, or perhaps suggest that the potential immunosuppressive functions of CGB genes are only one part of the mechanism underlying their association with poor prognosis. For example, hCG facilitates many pro-decidualization and pro-tumorigenic processes such as invasion, angiogenesis, in addition to immunosuppression. Our preliminary analyses suggest that these associations with *CGB7* are independent of *CGB3, CGB5* and *CGB8. CGB7* and bulk *CGB3, CGB5, and CGB8* are each independently associated with ICI resistance. Additional analyses to tease apart the isolated contributions of each gene will be required, and larger cohorts and study sizes will facilitate this. We hypothesize that, based on our findings in this study, while *CGB7* may uniquely be a marker of immune suppression and has great value as a marker or poor prognosis based on our analyses, that expression of any CGB gene has value as a marker of ICI resistance and worsened survival probability.

As primary and acquired resistance to checkpoint inhibitor immunotherapies remains a significant barrier to long-term remission for the majority of patients, further validation of *CGB7* as a biomarker and characterization of the mechanism by which *CGB7* suppresses the anti-cancer effects of ICIs may help us predict patient response, inform treatment, and perhaps even identify CGB7 as a potential therapeutic target. As CGB proteins are predominantly cancer-specific and readily detectable analytes in urine and plasma as the basis of modern pregnancy tests, this highlights the feasibility of CGB as a cancer biomarker. Further work will be required to validate CGB7 as a pan-cancer biomarker by confirming associations between *CGB7* expression and immunosuppressive signatures in other cancer types, and ultimately by validating this association between *CGB7* expression and ICI response in patient samples.

Ultimately, targeting CGB genes could present a promising approach for combination therapy with immunotherapies such as immune checkpoint inhibitors. As CGB proteins are secreted, there is substantial potential for development of an inhibitory anti-pan-CGB or anti-CGB7 antibody. There is precedent for potential therapeutic value of antibody neutralization of CGB proteins in the form of anti-cancer vaccines (Moulton 2002), and an anti-CGB therapeutic antibody could potentially provide therapeutic value on its own, or to provide specificity to a coupled anti-cancer payload.

## METHODS

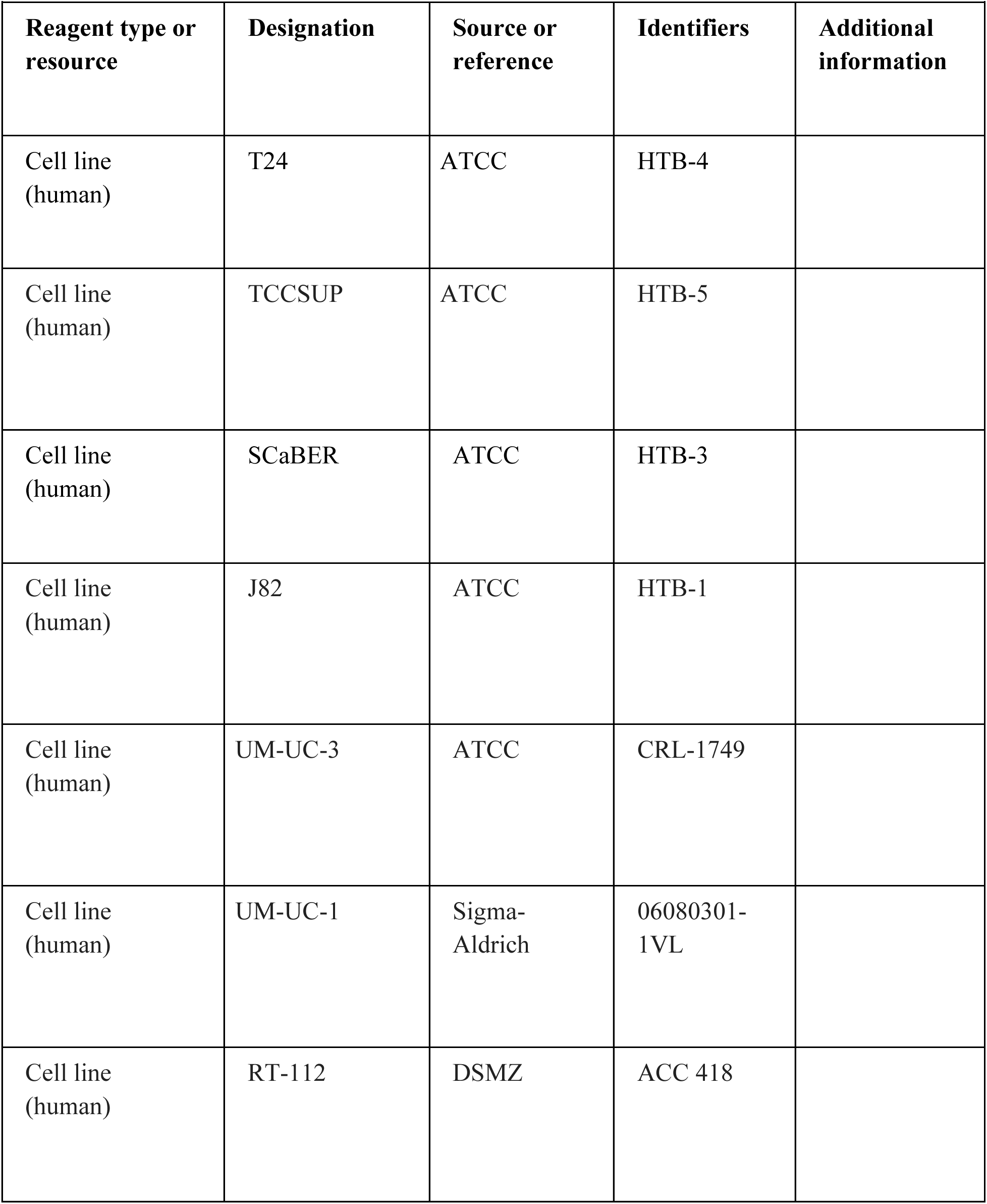
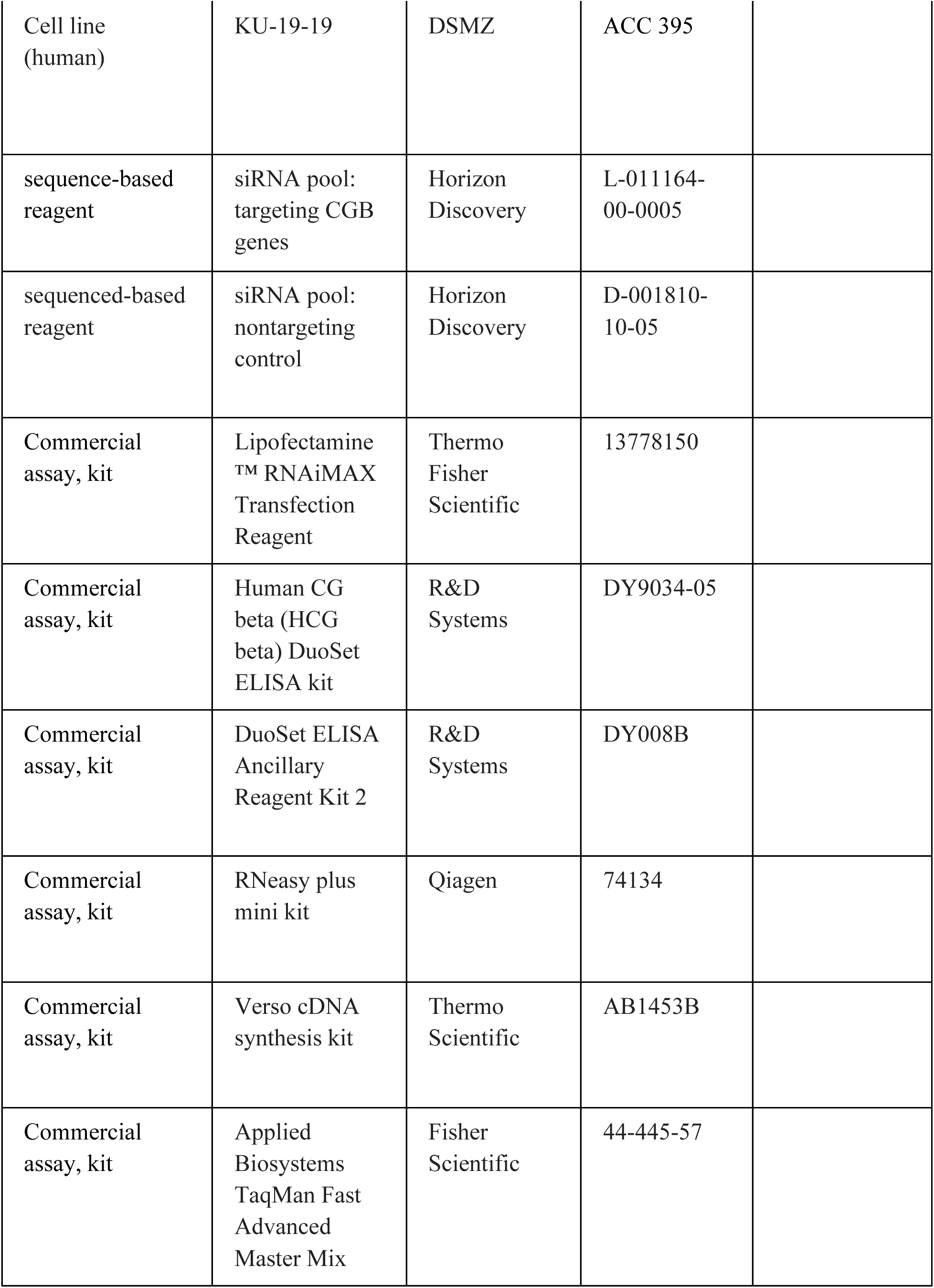
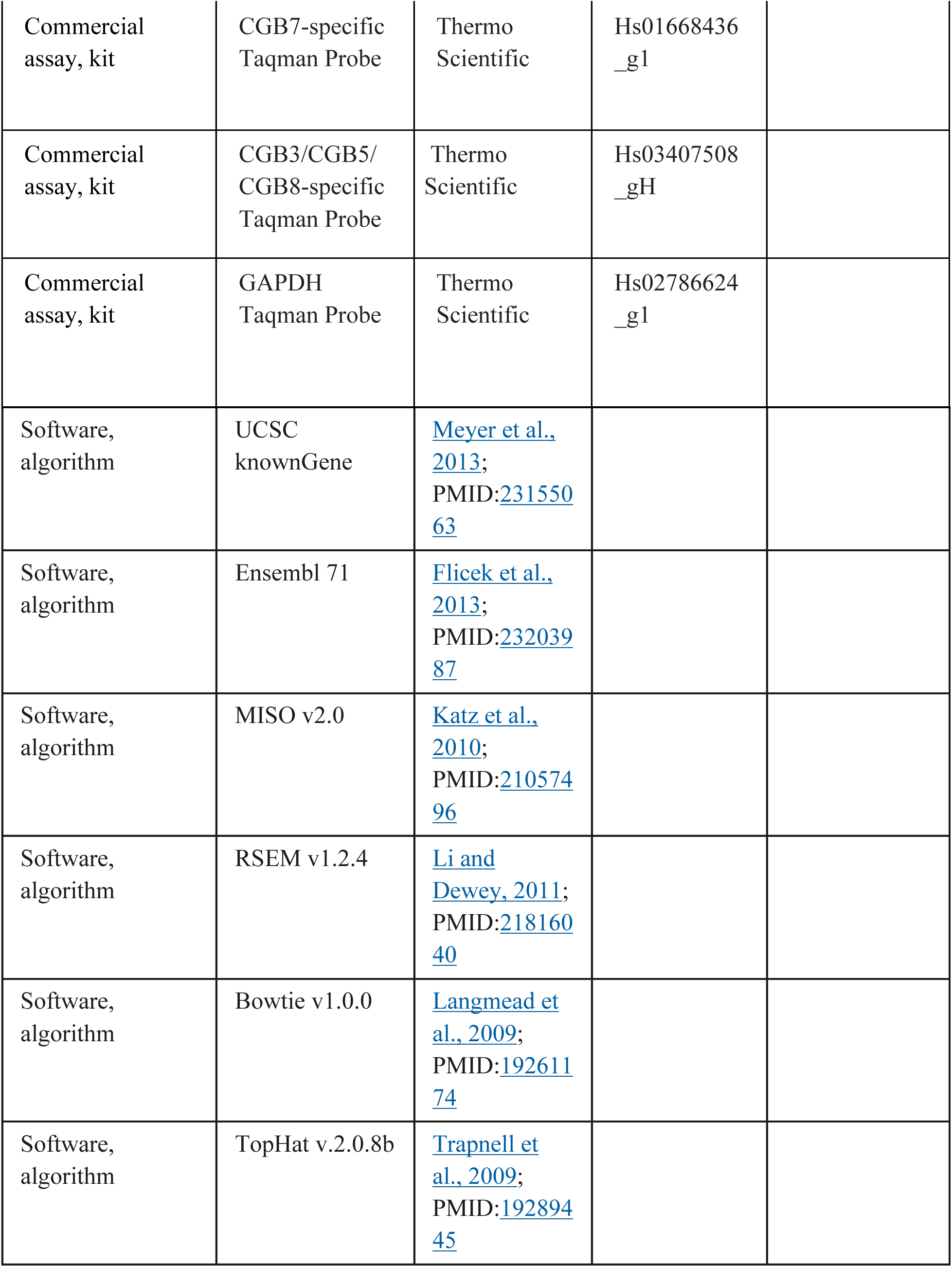
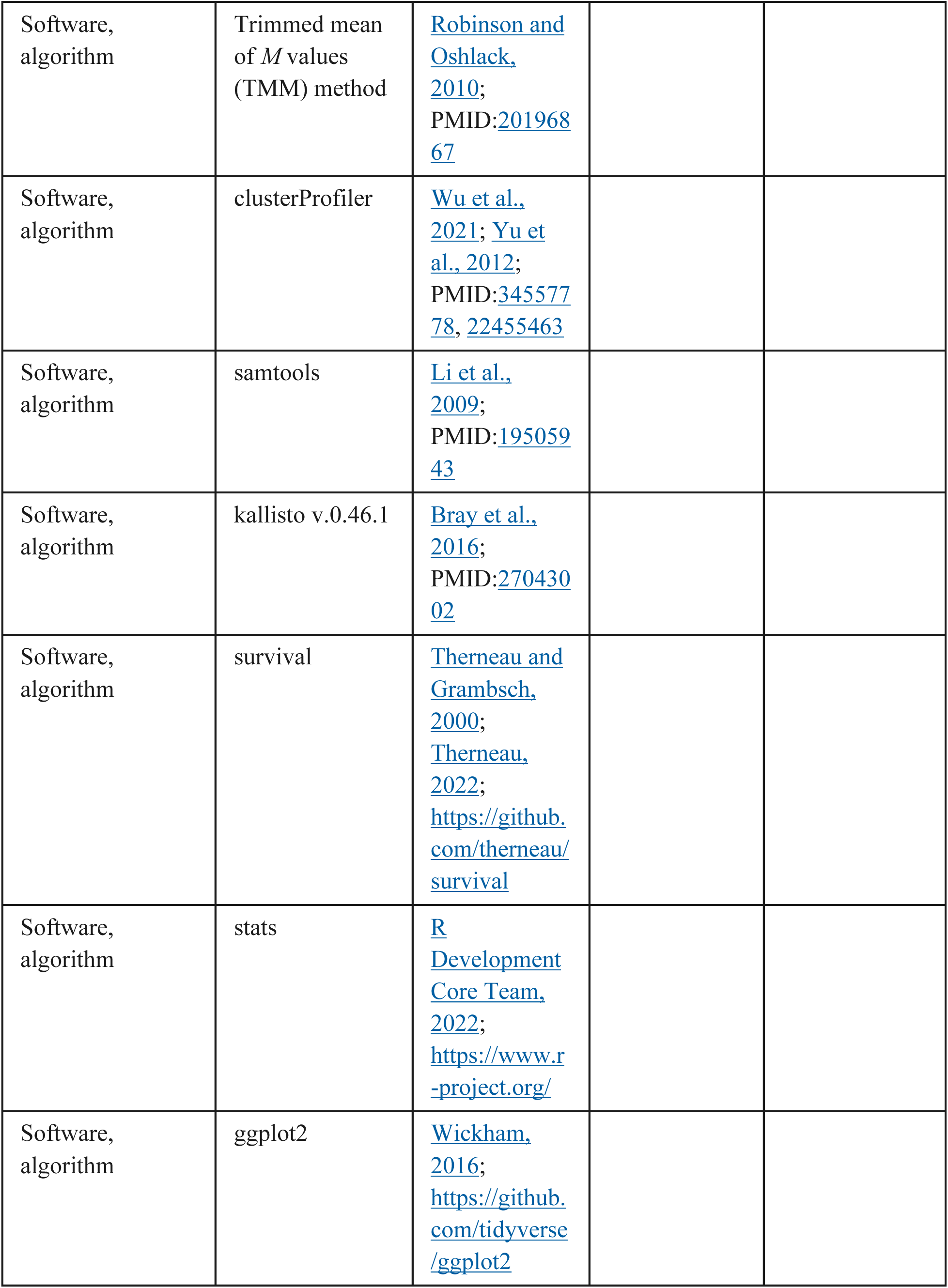
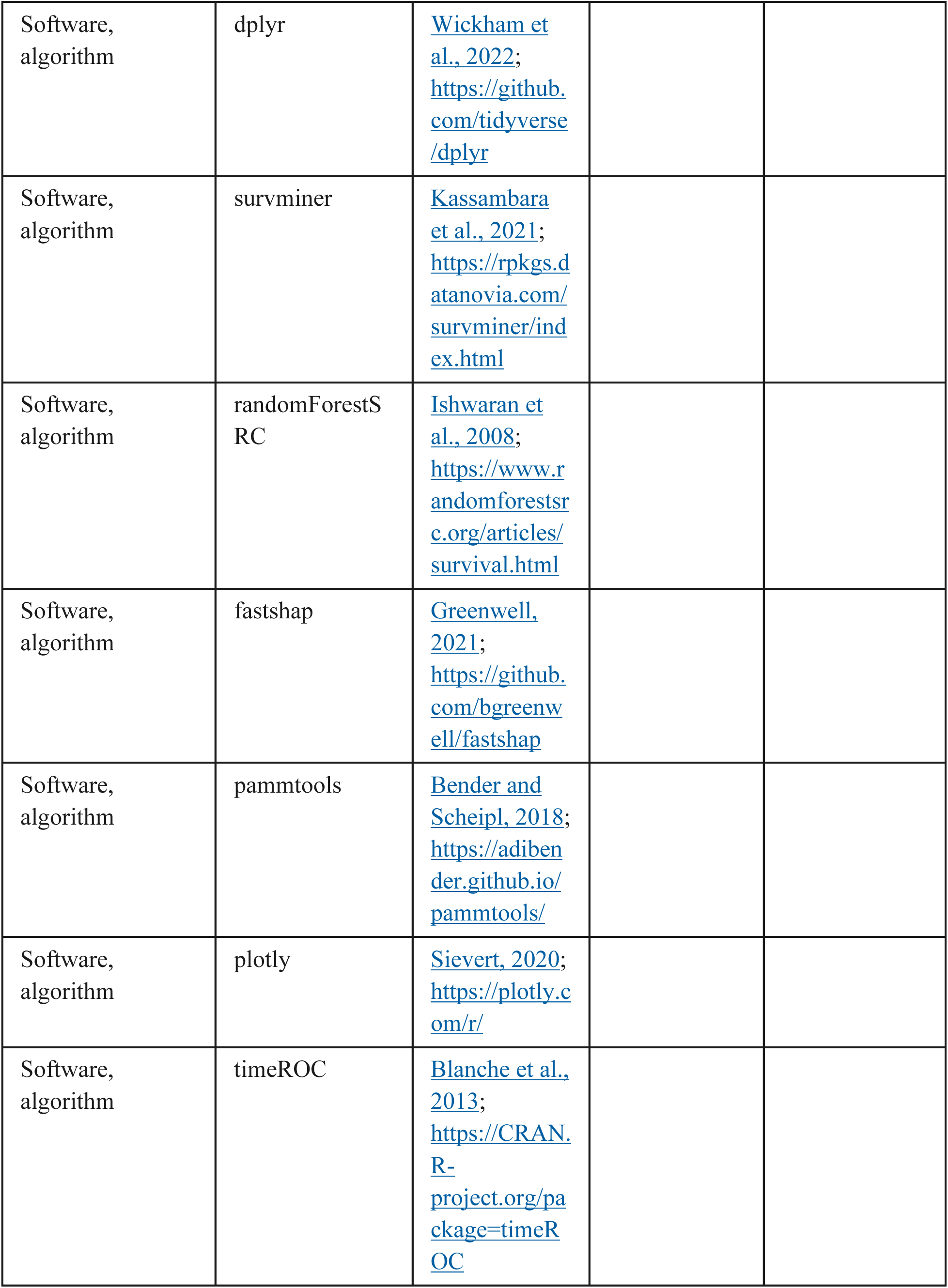
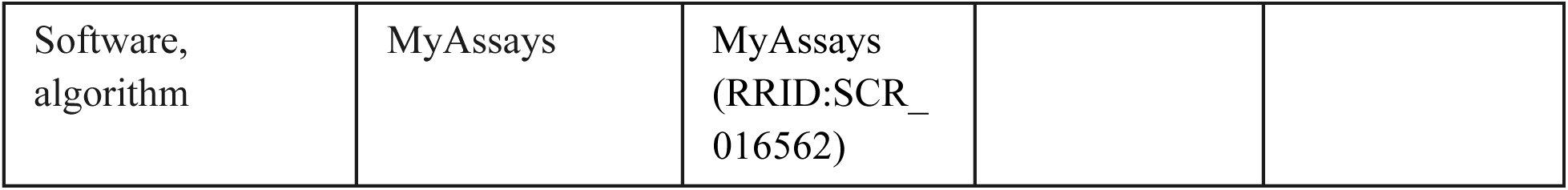
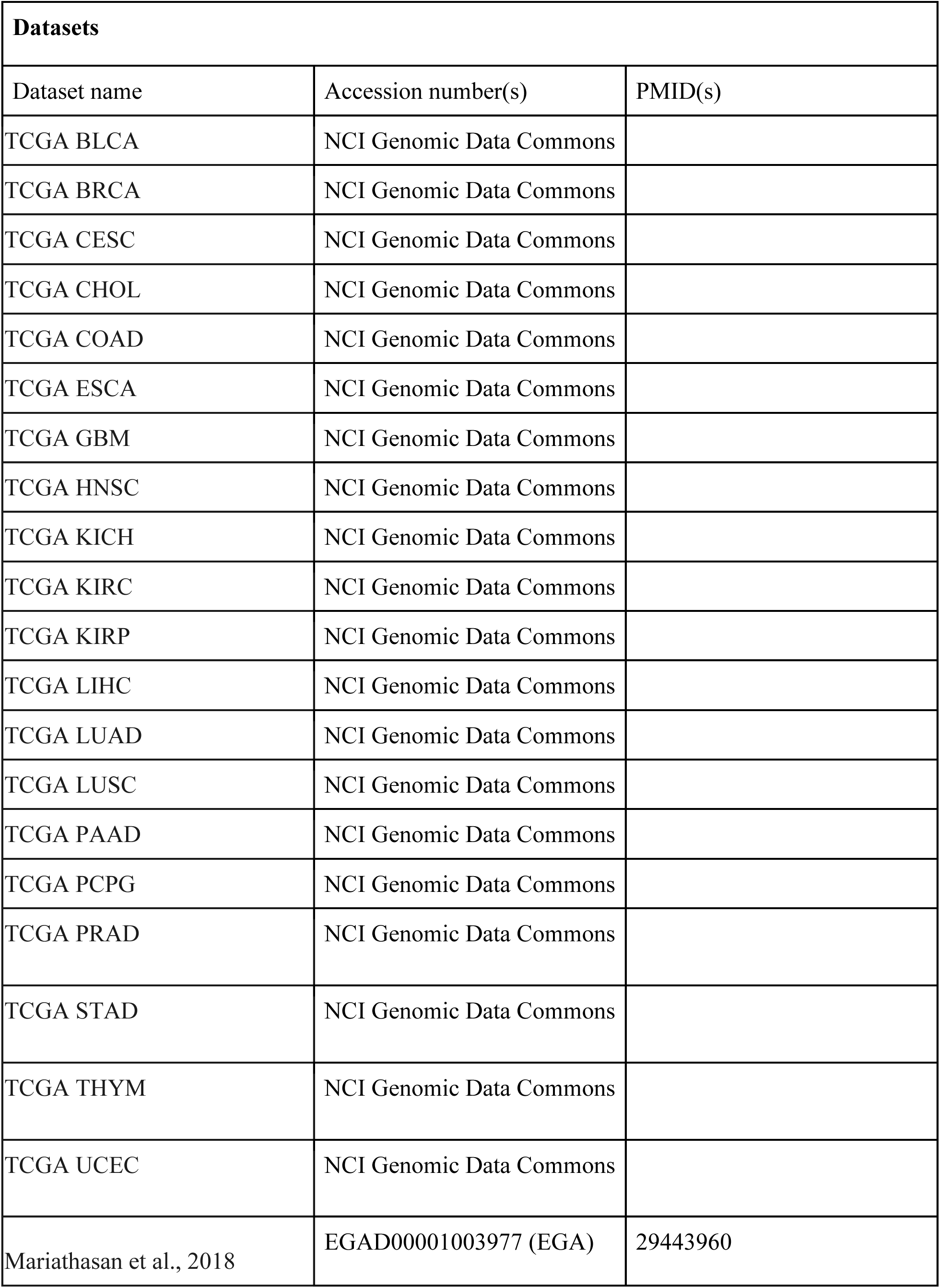

### Cell lines

All cell lines utilized are human urothelial cancer cell lines cultured at 37°C and 5% CO2. SCaBER, T24, TCCSUP, J82, UM-UC-3, cell lines were purchased from ATCC (ATCC, Manassas, VA). RT-112 and KU-19-19 cell lines were purchased from DSMZ (DSMZ, Braunschweig, Germany). UM-UC-1 cell line was purchased from Sigma-Aldrich (St. Louis, MO). SCaBER, RT-112, UM-UC-1, UM-UC-3, TCCSUP and J82 cells were cultured in Eagle’s Minimum Essential Medium (ATCC) supplemented with 10% FBS (Gibco, Waltham, MA) and 1% Pen Strep (Gibco). T24 cells were cultured in McCoy’s 5A medium supplemented with 10% FBS and 1% Pen Strep. KU-19-19 cells were cultured in RPMI 1640 (Gibco) medium supplemented with 10% FBS and 1% Pen Strep.

### CGB knockdown experiments

siRNA pools targeted all CGB genes (Horizon Discovery, Cambridge, Cambridgeshire, UK) or were a non-targeting control pool (Horizon Discovery). siRNAs transfected into cells of approximately 70% confluence with Lipofectamine™ RNAiMAX Transfection Reagent (Thermo Fisher Scientific, Waltham, MA) per the manufacturer’s instructions.

## ELISA

Conditioned medium: conditioned medium was collected from each indicated cell line after 72 hours of subculture in the medium described above. Bulk beta-hCG protein levels were measured and quantified using a Human CG beta (HCG beta) DuoSet ELISA kit (R&D Systems, Minneapolis, MN) DuoSet ELISA Ancillary Reagent Kit 2 (R&D Systems) according to the manufacturer’s instructions. Media was diluted 1:10 from all cell lines with the exception of KU-19-19, which was diluted 1:100 prior to assay input due to high CGB protein levels. Cells were counted at collection to approximate concentration of CGB per 10,000 cells. Experiments were run in biological and technical triplicate. Raw O.D. values of standards were fit to a 4PL curve using MyAssays software.

### qRT-PCR

Total RNA was extracted using a Qiagen RNeasy plus mini kit according to the manufacturer’s instructions (Qiagen, Hilden, Germany). RNA was reverse transcribed to cDNA using a Verso cDNA synthesis kit (Thermo Scientific). *CGB7* transcripts detected with Applied Biosystems TaqMan Fast Advanced Master Mix (Fisher Scientific), *CGB7*-specific Taqman Probe (Thermo Scientific), *CGB3*/*CGB5*/*CGB8*-specific Taqman Probe (Thermo Scientific), and GAPDH Taqman Probe (Thermo Scientific). Experiments were run in biological and technical triplicate. Experiments were run on a ABI QuantStudio 5 Real-Time PCR System (Thermo Fisher Scientific). Data was analyzed using the ΔΔCt method.

### RNA sequencing analysis, genome annotations and differential gene expression

RNA-sequencing data was analyzed as previously described (Pineda 2024). RNA-seq reads were mapped to an annotated transcriptome created using Ensembl 71 (Flicek et al., 2013), UCSC knownGene (Meyer et al., 2013), and MISO v2.0 (Katz et al., 2010) annotations for the hg19/GRCh37 assembly using RSEM version 1.2.4 (B. Li & Dewey, 2011) modified to call Bowtie v1.0.0 with option ‘-v 2’ (Langmead et al., 2009). Unaligned reads were then mapped to the hg19/GRCh37 genome assembly using TopHat version 20.8b (Trapnell et al., 2009). Gene expression values in TPM (transcripts per million) were normalized via the trimmed mean of M values (TMM) method (M. D. Robinson & Oshlack, 2010). For TCGA studies, we analyzed all available samples across 20 distinct cancer types that included at least one matched non-tumor tissue sample. Differential gene expression to compare positive and negative samples for expression of CGB genes in both the TCGA datasets and the dataset from the IMvigor210 clinical trial cohort (Mariathasan et al. 2018, Balar et al., 2017) was calculated via Mann-Whitney U Test and using several thresholds: the absolute value of log-fold-change threshold of 1.5 or greater, a maximum p-value of 0.05, a minimum bayes factor of 100, and maximum FDR of 0.01. We defined positive expression of CGB genes as TPM >1 and negative expression as TPM < 0.25 unless otherwise noted.

### Clinical variable analyses

Clinical variables were reported for patients enrolled in the IMVigor210 clinical trial (Mariathasan et al., 2018, Balar et al., 2017). Patient response to Atezolizumab was scored according to the Response Evaluation Criteria for Solid Tumors (RECIST) (Balar et al., 2017). Tumor subtype was determined according to Lund 2 taxonomy for bladder cancer classification (Mariathasan 2018). Immunophenotype was determined based on the level and pattern of CD8+ T cell infiltration by CD8a IHC staining (Mariathasan et al., 2018). For each analysis, patients were first stratified by expression of each CGB gene, or by summed expression of multiple CGBs where indicated. NAs were filtered out for each response, immunophenotype, and subtype analysis individually after CGB gene stratification. We determined the proportions of each RECIST score, immunophenotype, and subtype in CGB-high (>75%) and CGB-low/negative (<25%) tumors, and quantified the difference in proportions via 1-sample proportions test with continuity correction.

### Survival analyses

Survival analyses were performed with the Kaplan–Meier estimator and statistical tests were performed with a logrank test (R package survival) (Therneau 2024). Samples were stratified by expression of individual genes, where positive expression was defined as exhibiting TPM >1, and negative expression was defined as exhibiting TPM <0.25. Adjusted survival probabilities were calculated by fitting a Cox Proportional Hazards model to adjust for the covariates sex and tumor mutational burden.

### Random Survival Forest and Feature Importance computation

The Random Survival Forest Model, Shapley value estimation, and partial dependence analyses was implemented as previously described (Pineda 2024). In brief, we randomly assigned patients into training (70%) and test (30%) datasets. We determined optimal hyperparameters via a grid search. That is, we evaluated 10,608 RSF models representing various values and combinations of the RSF hyperparameters: number of trees, minimum terminal node size, number of randomly selected splitting variables, handling of missing data, splitting rule, and bootstrapping method.

The model which minimized both the OOB training and the test errors (defined as 1 − concordance index) was selected: ntree = 1000, nodesize = 6, mtry = 3, na.action = “na.impute”, splitrule = “logrank”, and samptype = “swr”. We set the hyperparameter nsplit = 0 to evaluate all possible split points. The predictions associated with the test cohort were handled using na.action = “na.omit” which excluded patients with missing data. Shapley values were calculated using the fastshap package (Greenwell, 2024). We used 1000 Monte Carlo repetitions and set the parameter adjust = TRUE to correct the estimated Shapley values such that local accuracy was satisfied. Shapley values associated with mortality and per time point overall survival predictions from the RSF model were used to quantify overall feature importance and time-dependent importance, respectively. The marginal effect of CGB7 expression on survival probability was assessed using partial dependence, implemented using the partial() function from the randomForestSRC package. Visualizations were created in the R programming environment using the dplyr (Wickham et al., 2022), ggplot2 (Wickham, 2016), pammtools (Bender and Scheipl, 2018), and plotly (Sievert, 2020) packages.

### Measuring survival model predictive accuracy

We evaluated the RSF model’s accuracy over time using time-dependent ROC curve analyses. For each timepoint, we calculated the cumulative/dynamic area under the ROC curve (AUC^C/D^) and 95% confidence interval using the timeROC package, which additionally corrects for bias due to right-censoring (Blanche et al., 2013). The training (OOB) or the test predictions for mortality were used as input.

## ACKNOWLEDGEMENTS

R.K.B. was supported in part by the NIH/NCI (R01 CA251138), NIH/NHLBI (R01 HL128239, R01 HL151651), and the Blood Cancer Discoveries Grant program through the Leukemia & Lymphoma Society, Mark Foundation for Cancer Research, and Paul G. Allen Frontiers Group (8023-20). R.K.B is a Scholar of The Leukemia & Lymphoma Society (1344-18) and holds the McIlwain Family Endowed Chair in Data Science. The results shown here are in part based upon data generated by the TCGA Research Network: https://cancergenome.nih.gov/. This research was supported by the Genomics Shared Resource of the Fred Hutch/University of Washington/Seattle Children’s Cancer Consortium (P30 CA015704). Figure 1A Created in BioRender.com.

## AUTHOR CONTRIBUTIONS

S.A.M. and R.K.B. designed the study and wrote the manuscript. S.A.M and J.M.B.P. executed and optimized experiments, performed analyses, and interpreted data. J.M.B.P. contributed to the writing of the manuscript. S.X.L, R. L., and A.S.C. conducted experiments and provided interpretation. S.X.L and E.W.N. contributed to project design.

---

**Supplemental Figure 1:**
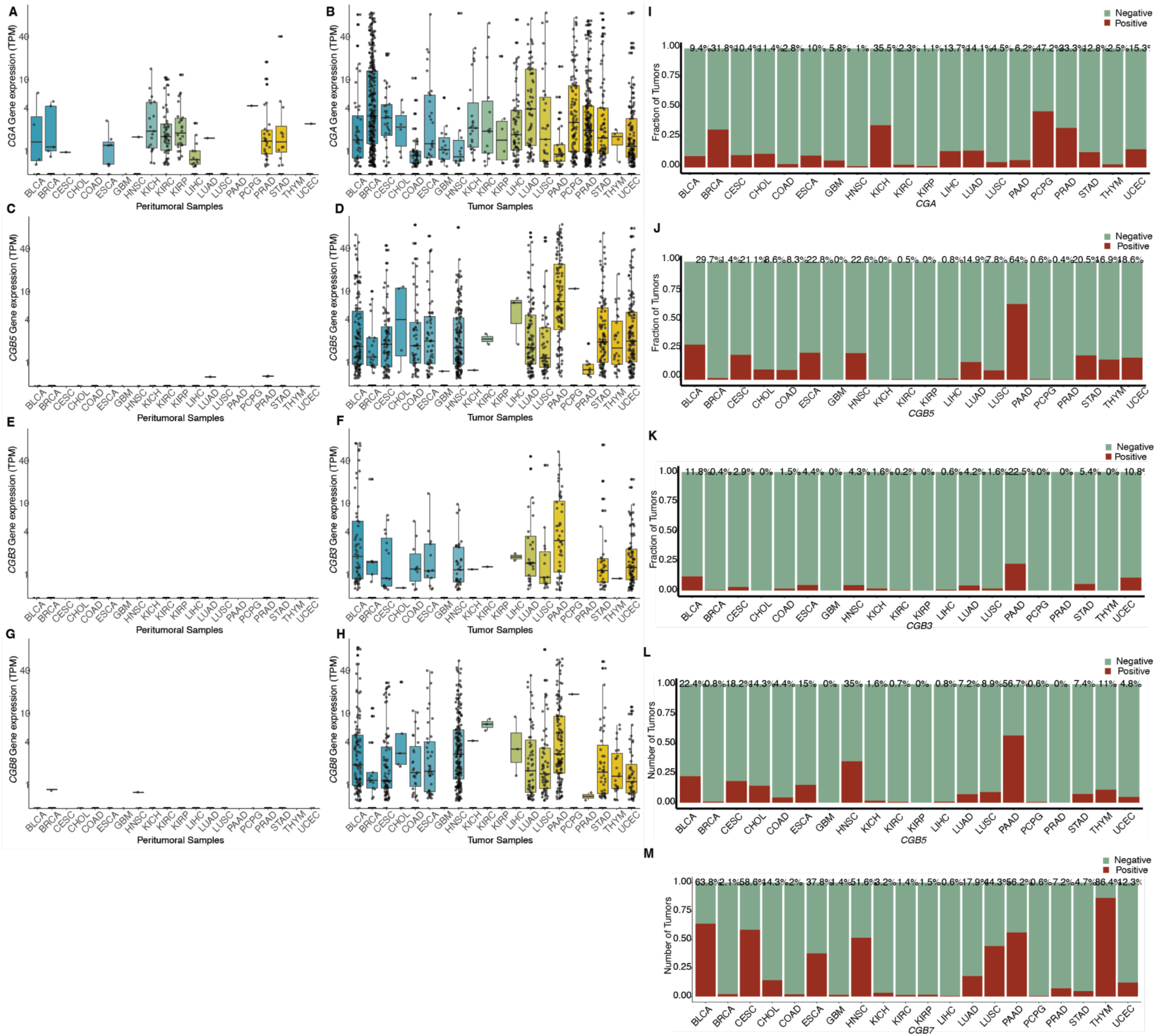
**CGB is expressed in multiple cancer types.** *CGA* mRNA expression in healthy peritumoral tissue samples (**A**) and matched tumor tissue samples (**B**) across 20 cancer types from the TCGA. Cancer type reflects the site of the primary tumor. *CGB3* mRNA expression in healthy peritumoral tissue samples (**C**) and matched tumor tissue samples (**D**) across 20 cancer types from the TCGA. Cancer type reflects the site of the primary tumor. *CGB5* mRNA expression in healthy peritumoral tissue samples (**E**) and matched tumor tissue samples (**F**) across 20 cancer types from the TCGA. Cancer type reflects the site of the primary tumor. *CGB8* mRNA expression in healthy peritumoral tissue samples (**G**) and matched tumor tissue samples (**H**) across 20 cancer types from the TCGA. Cancer type reflects the site of the primary tumor. (**I, J, K, L, M**) Fraction of tumors across 20 cancer types expressing *CGA*, *CGB3*, *CGB5*, *CGB8*, or *CGB7*, gene expression with TPM >1, respectively. Percent positive is annotated above each cancer type. TPM = transcripts per million.

**Supplemental Figure 2:**
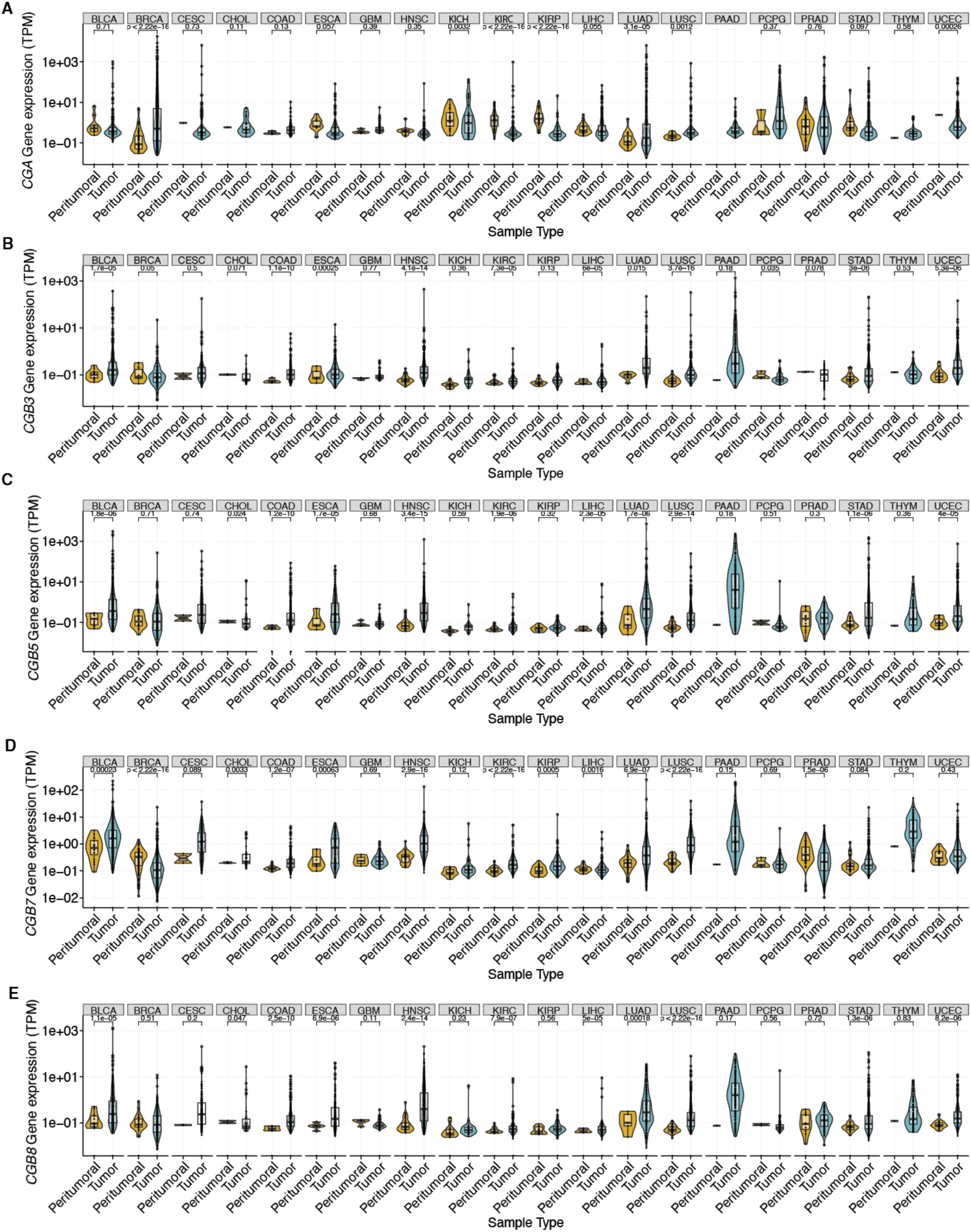
**CGB expression is upregulated across cancer types.** Violin plots of *CGA* (**A**), *CGB3* (**B**), *CGB5* (**C**), *CGB7* (**D**), or *CGB8* (**E**) expression in tumors and matched healthy peritumoral tissue samples across cancer type datasets from The Cancer Genome Atlas. Expression is in transcripts per million (TPM). P-values determined by wilcox signed rank test.

**Supplemental Figure 3:**
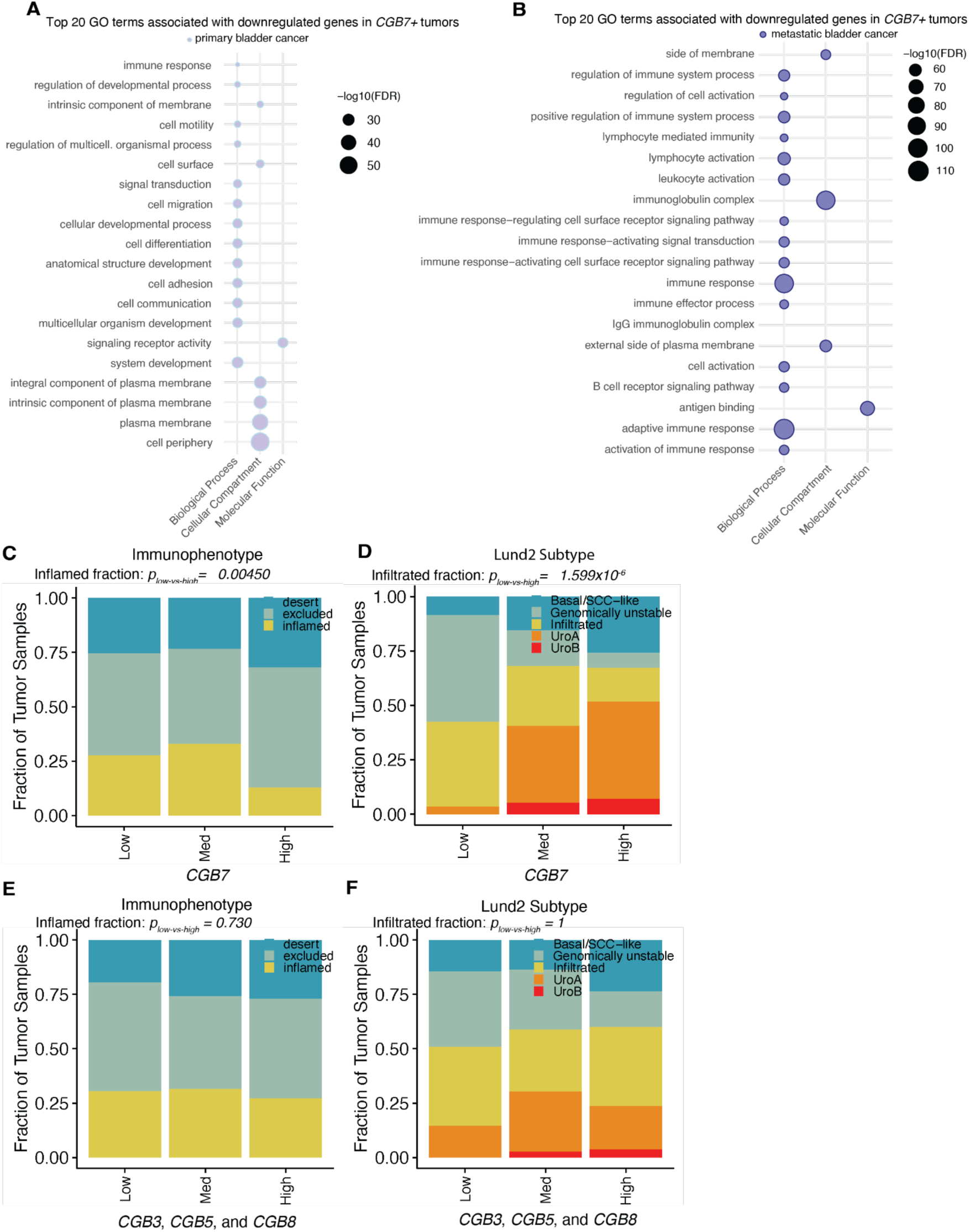
**CGB expression is associated with altered immune infiltrate.** (**A**) The top 20 Gene Ontology terms associated with *CGB7* expression in primary urothelial cancers in the TCGA BLCA dataset. (**B**) The top 20 Gene Ontology terms associated with *CGB7* expression in metastatic urothelial cancers in the IMVigor 210 clinical trial dataset (data re-analyzed from Mariathasan et al., 2018). (**C**) Immunophenotype data: immune desert, immune excluded, or inflamed. Tumors with expression of *CGB3*, *CGB5*, or *CGB8* > 1 TPM were removed, and remaining samples stratified by *CGB7* expression. P-values determined by proportions test with continuity correction. Chi-squared = 8.072, df = 1, p-value = 0.0045. (**D**) Tumor subtype data: basal/SCC-like, genomically unstable, immune infiltrated, urothelial type A, or urothelial type B. Tumors with expression of *CGB3*, *CGB5*, or *CGB8* > 1 TPM were removed, and remaining samples stratified by *CGB7* expression. P-values determined by proportions test with continuity correction. Chi-squared = 23.025, df = 1, p-value = 1.60×10^-6^. (**E**) Immunophenotype data: immune desert, immune excluded, or inflamed. Tumors with expression of *CGB3*, *CGB5*, or *CGB8* > 1 TPM were removed, and remaining samples stratified by *CGB7* expression. P-values determined by proportions test with continuity correction. Chi-squared = 0.119, df = 1, p-value = 0.730. (**F**) Tumor subtype data: basal/SCC-like, genomically unstable, immune infiltrated, urothelial type A, or urothelial type B. Tumors with expression of *CGB7* > 1 TPM were removed, and remaining samples stratified by summed *CGB3*, *CGB5*, and *CGB8* expression. P-values determined by proportions test with continuity correction. Chi-squared = 0, df = 1, p-value = 1. Med = medium.

**Figure S4:**
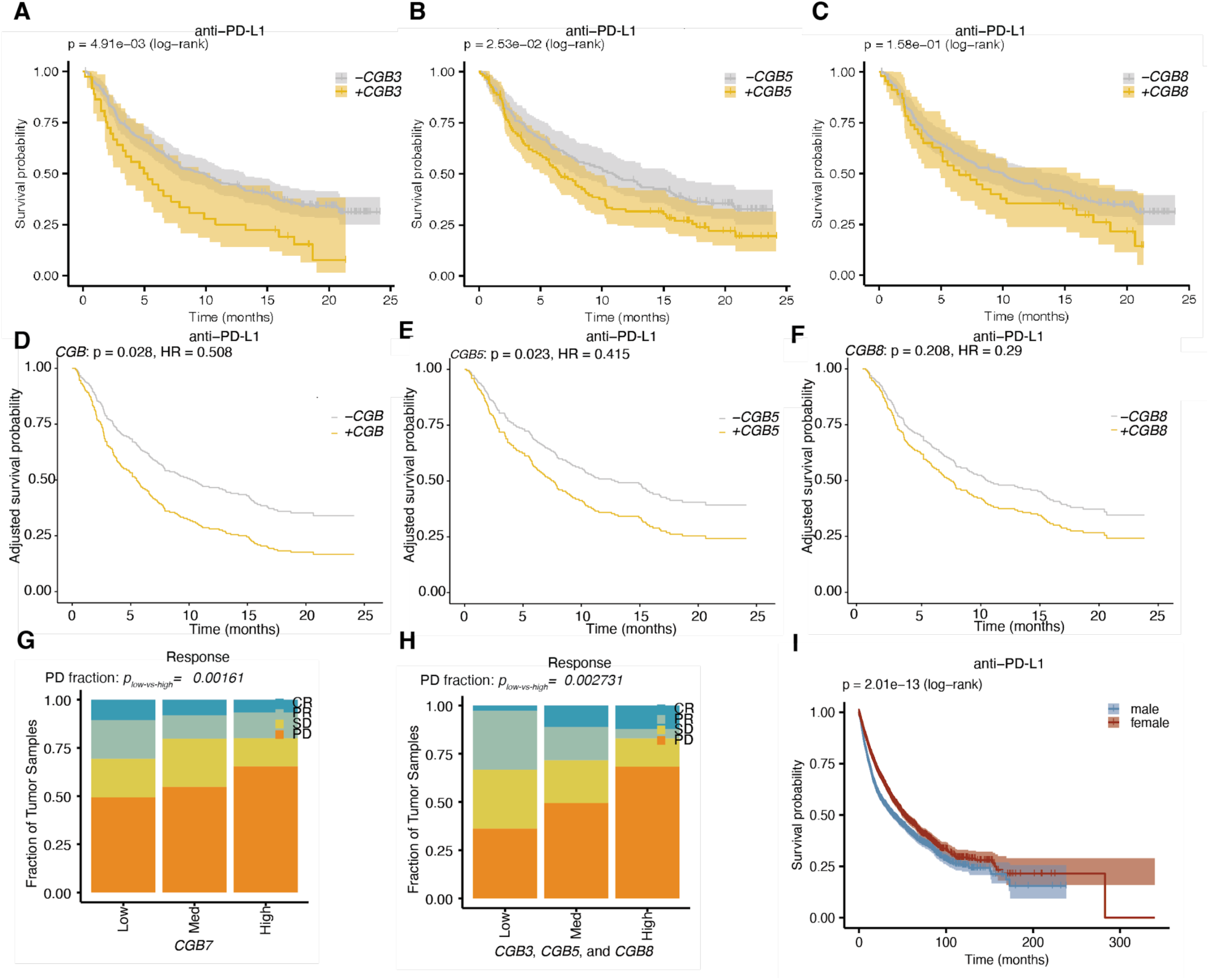
**CGB expression is associated with decreased response to ICI therapy.** (**A**) Kaplan Meier overall survival curves comparing patients with *CGB3*+ (gold) and *CGB3*-(gray) advanced urothelial cancer tumors (data re-analyzed from Mariathasan et al., 2018). All patients received Atezolizumab (anti-PD-L1). P-value determined by log rank test. (**B**) As in (**A**), but comparing patients with *CGB5* positive (gold) and negative (gray) tumors. (**C**) As in (**A**), but comparing patients with *CGB8* positive (gold) and negative (gray) tumors. (**D**) Kaplan Meier overall survival curves as shown in (**A**) adjusted for confounding effects of sex and tumor mutational burden covariates by cox proportional hazards modeling. Hazard ratio (HR) and p-value obtained from fitting a cox proportional hazards model. (**E**) Kaplan Meier overall survival curves as shown in (**B**) adjusted as described in (**D**). (**F**) Kaplan Meier overall survival curves as shown in (**C**) adjusted as described in (**D**). (**G**) Response determined by RECIST (Mariathasan 2018). Tumors with expression of *CGB3*, *CGB5*, or *CGB8* > 1 TPM were removed, and remaining samples stratified by *CGB7* expression. P-values determined by proportions test with continuity correction. Chi-squared = 9.954, df = 1, p-value = 0.00161. (**H**) Response determined by RECIST (Mariathasan et al., 2018). Tumors with expression of *CGB7* > 1 TPM were removed, and remaining samples stratified by summed *CGB3*, *CGB5*, and *CGB8* expression. P-values determined by proportions test with continuity correction. Chi-squared = 8.979, df = 1, p-value = 0.00273. (**I**) Kaplan Meier overall survival curves comparing male (blue) and female (red) patients with advanced urothelial carcinoma tumors. All patients received Atezolizumab (anti-PD-L1). P-value determined by log rank test. Response Evaluation Criteria in Solid Tumors (RECIST) scoring: CR = complete response, PR = partial response, SD = stable disease, PD = progressive disease. Med = medium.

**Figure S5:**
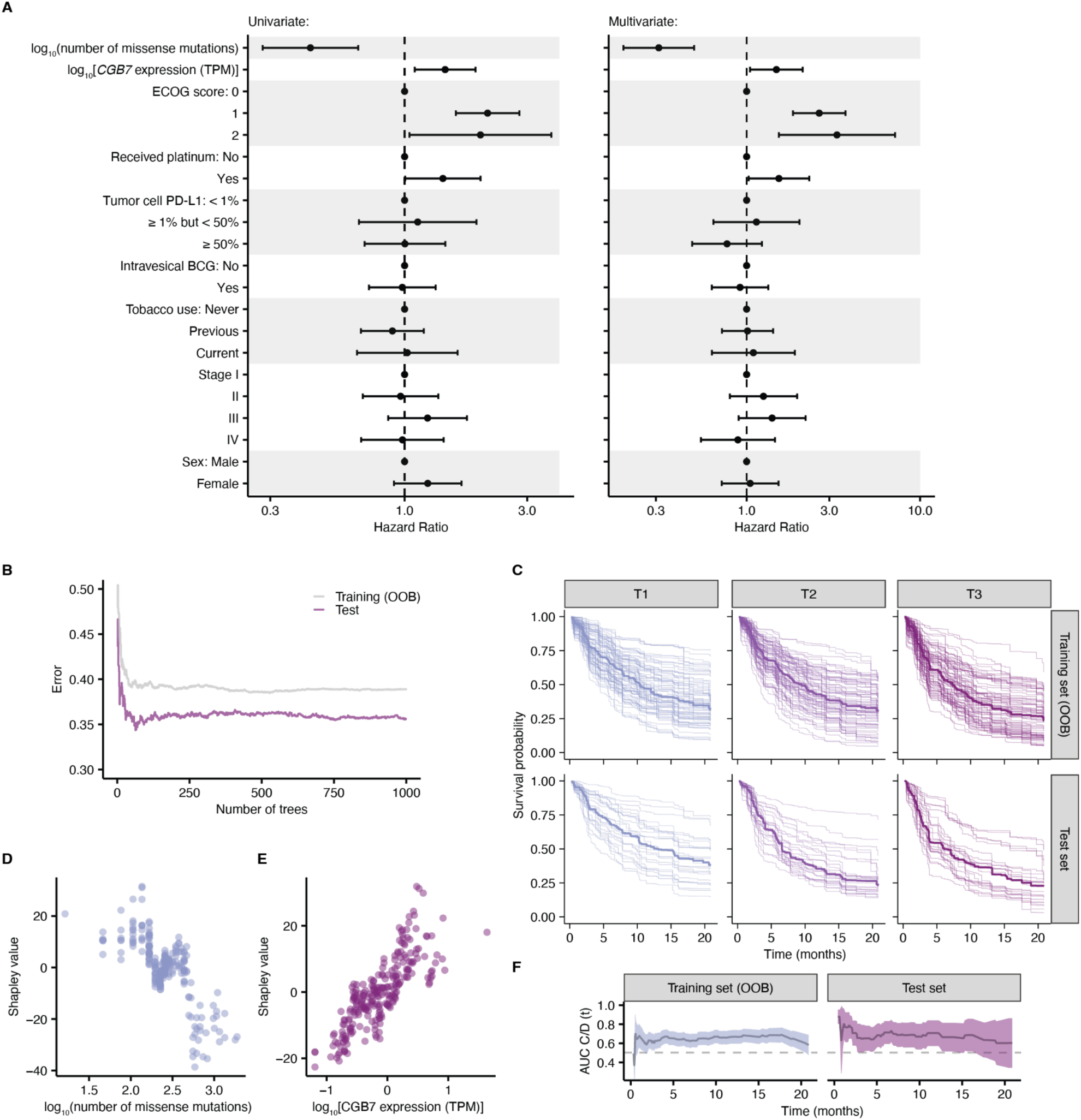
**Statistical and machine learning models exhibit the negative effects of *CGB7* expression on overall survival.** (**A**) Hazard ratios estimated by Cox Proportional Hazards Regression in the univariate (left) and multivariate (right) contexts. Error bars denote the 95% confidence interval of the hazard ratio. (**B**) The training out-of-bag error (OOB error, solid gray line) and the test error (solid purple line) as a function of the number of trees in the Random Survival Forest (RSF) model. Error is defined as 1 − Harrell’s concordance index. (**C**) RSF predicted overall survival for individual patients (thin lines) stratified into terciles, by *CGB7* expression. OOB survival predictions are shown for the patients in the training set. The median survival function across the cohort is shown (thick line). (**D**) Shapley dependence plot correlating tumor mutational burden (TMB, number of missense mutations) and RSF mortality. Each patient is represented by a single point. (**E**) As in (**D**), but showing *CGB7* expression. (**F**) Time-dependent receiver operating characteristic (ROC) analyses. The cumulative/dynamic area under the ROC curve (AUC^C/D^) (solid line) and associated 95% confidence interval (transparent ribbon) are calculated for the training and test sets. The training set AUC^C/D^ was calculated using the RSF OOB mortality predictions.

